# The evolution of signalling and monitoring in plant-fungal networks

**DOI:** 10.1101/2024.08.15.608109

**Authors:** Thomas W. Scott, E. Toby Kiers, Stuart A. West

**Affiliations:** Department of Biology, University of Oxford, 11a Mansfield Road, Oxford, United Kingdom; School of Biology, University of St Andrews, Dyers Brae, St Andrews, United Kingdom; A-life, Vrije Universiteit Amsterdam, Amsterdam, the Netherlands; Society for the Protection of Underground Networks, SPUN, Dover, DE 19901, USA

## Abstract

Experiments have shown that when one plant is attacked by a pathogen or herbivore, this can lead to other plants connected to the same mycorrhizal network upregulating their defence mechanisms. It has been hypothesised that this represents signalling, with attacked plants producing a signal to warn other plants of impending harm. We examined the evolutionary plausibility of this and other hypotheses theoretically. We found that the evolution of plant signalling about an attack requires restrictive conditions, and so will rarely be evolutionarily stable. The problem is that signalling about an attack provides a benefit to competing neighbours, even if they are kin, and so reduces the relative fitness of signalling plants. Indeed, selection is often more likely to push plant behaviour in the opposite direction – with plants signalling dishonestly about an attack that has not occurred, or by suppressing a cue that they have been attacked. Instead, we show that there are two viable alternatives that could explain the empirical data: (1) the process of being attacked leads to a cue (information about the attack) which is too costly for the attacked plant to fully suppress; (2) mycorrhizal fungi monitor their host plants, detect when they are attacked, and then the fungi signal this information to warn other plants in their network. Our results suggest the empirical work that would be required to distinguish between these possibilities.

**Significance statement:** Experiments have shown that when one plant is attacked by a herbivore, this can lead to other plants connected to the same mycorrhizal network upregulating their defence mechanisms. It has been hypothesised that this represents signalling, with attacked plants producing a signal to warn other plants of impending harm. We found theoretically that plant warning signals are rarely evolutionarily stable. Instead, we identify two viable alternatives that could explain the empirical data: (1) being attacked leads to a cue (information about the attack) which is too costly for the attacked plant to suppress; (2) mycorrhizal fungi monitor their host plants, detect when they are attacked, and then the fungi signal this information to warn other plants in their network.

## Introduction

Mycorrhizal fungi form symbiotic associations with plant roots, trading nutrients such as phosphorous and nitrogen for plant-derived carbon. These fungi form physical networks of mycelium that can connect roots of different plants and act as potential routes for signalling between those plants (1–7). Several laboratory experiments have provided clear evidence that when one plant in a mycorrhizal network is attacked that this leads to other plants in the network upregulating their defence mechanisms. For example, when a tomato plant is infested with a leaf chewing caterpillar, tomato plants connected to the same network will increase their production of defence enzymes (8). It has been hypothesised that this pattern represents a ‘warning signal’ in which the attacked plant actively signals to other plants using chemicals transported via the mycorrhizal network. This work has even fuelled narratives in the media that forest trees use mycorrhizal networks to warn other trees of impending danger (9, 10).

However, the evolutionary plausibility of this signalling hypothesis remains unclear (2, 11–24). A signal is defined as ‘*any act or structure that alters the behaviour of other organisms, which evolved owing to that effect, and which is effective because the receiver’s response has also evolved*’ (25, 26). Consequently, signalling is a form of cooperation, that is only favoured when it provides a benefit to both the sender and the receiver (25). Plants compete with neighbours for resources such as sunlight and nutrients, and so helping a neighbour could be costly to a potential signaller (27). Neighbouring plants could be relatives, which could provide a kin-selected benefit of signalling, but competition between relatives can reduce or even negate any benefit of helping relatives (28–31). For instance, a low migration rate could cause relatives to signal to each other, leading to a kin-selected benefit, but it could also cause relatives to compete with each other, negating this benefit. Consequently, it is not clear if signalling about attack would be evolutionarily stable between neighbouring plants who are both relatives and competitors.

In addition, there are at least two alternatives to plant signalling that could possibly explain the experimental data (Fig. 1) (32). One possibility is that the neighbouring plants could be detecting a cue that another plant is being attacked (27). A cue is defined by when a receiver uses some feature of the sender to guide their own behaviour, but this feature has not evolved for that purpose (26). An example of a cue is when a mosquito searching for a mammal to bite will fly up wind if it detects carbon dioxide (26). Mosquitos use carbon as a cue of the presence of a source of blood, but mammals do not produce carbon dioxide to signal their presence to mosquitoes (getting bitten is costly!). All that is required for a cue of plant attack is that the damage caused by attack causes the attacked plant to produce something, such as a released chemical (volatile) (33). The production of cues of herbivore-attack may be unavoidable (34).

**Figure 1.**
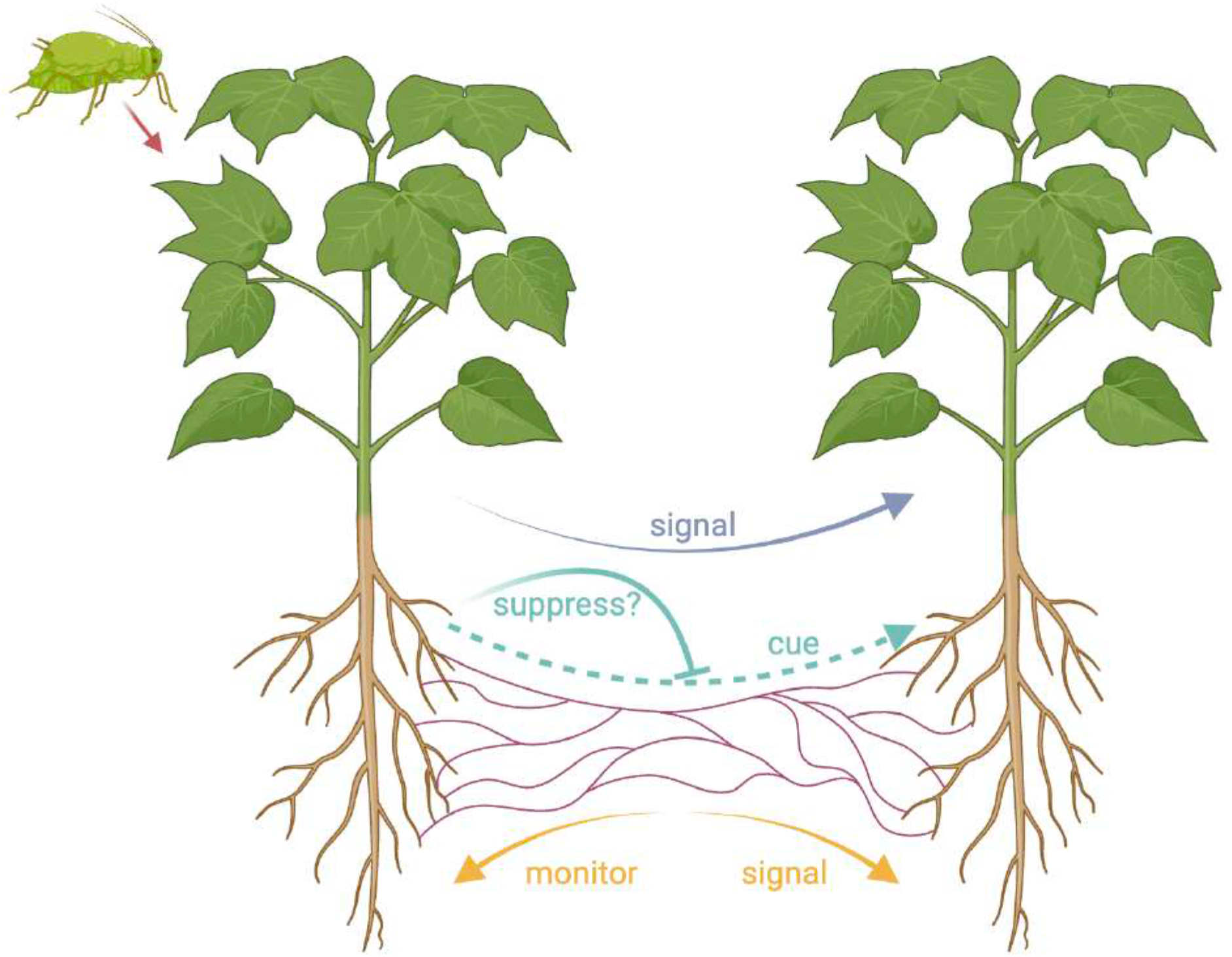
Hypotheses for information transfer in plant-fungal networks. Two plants are connected *via* a mycorrhizal network. One plant is under attack (*e*.*g*. by aphids), and this information may be transferred *via* the mycorrhizal network to the other plant, allowing it to upregulate its defence mechanisms. There are three hypotheses regarding how this information is transferred: (*blue*) signalling by the attacked plant; (*cyan*) cues, which are potentially vulnerable to suppression by the attacked plant; (*orange*) monitoring and signalling by the mycorrhizal network. This figure was made with https://Biorender.com.

A second possible alternative explanation of the experimental data is that the fungus can detect when a plant is being attacked, and that it then signals this information to the other plants to which it is connected (13). This would therefore involve fungi ‘monitoring’ plants to become aware of cues of herbivore attack, and it would be the fungus - not the attacked plant - that is signalling. Fungi could be selected to monitor and signal to plants in their network because they actively trade resources with those individuals, and so could benefit from helping keep them in better condition, to make better trading partners (27, 35, 36).

The difference among these hypotheses matters because they represent different evolutionary outcomes, imply that information transfer is favoured for different reasons, and require different empirical tests (26, 37, 38). With plant signalling, the problem is to determine how both the sender and receiver benefit. What conditions would be required for honest signalling to be evolutionary stable, and how could this seemingly altruistic act of warning neighbours be explained in the context of plant-plant competition? Could plants even be favoured to signal dishonestly to harm competitors? In contrast, with a cue, we would need to ask: what stimulates the cue to be produced? If the cue provides a benefit to neighbours, then does this cue impose a cost to the producer, by aiding their competitors? Why don’t plants suppress a cue? Finally, in the case of mycorrhizal monitoring, we would need to ask: why would the fungus be selected to monitor and then produce a signal in response? And what exactly is being monitored?

We investigated these different hypotheses theoretically, by applying kin selection and signalling theory to the question of herbivory information transfer between plants. Our aim was to determine their evolutionary plausibility, as well as define the empirical work required to test among them. We first consider selection on plants, examining whether: (a) an attacked plant would be selected to signal that it had been attacked; (b) plants can be selected to signal dishonestly about attack, to harm competitors; (c) plants can be selected to suppress a cue that they are being attacked. We then examine selection on mycorrhizal fungi and ask whether they can be selected to monitor plants, to determine whether they are being attacked, and then signal that information to neighbouring plants.

## Results

We constructed a series of theoretical models to examine selection on signals and cues, from the perspective of both plants and mycorrhizal fungi. We constructed deliberately simple models which are easy to interpret and can be applied across diverse species (39–41).

### Plant signalling model

We first examined whether an attacked plant can be selected to produce a warning signal. We assume an infinite population of individual plants, split into patches (demes) of size *N* (infinite island model) (42). Each generation, with probability *p*, the population is attacked, for instance by a herbivore or pathogen (with probability 1-*p*, the population is not attacked). In generations where the population is attacked, a random individual on each patch, *i*, is initially attacked, and suffers a fecundity cost of *d*. This individual then invests *x*_*i*_ into the production of a signal, resulting in an additional fecundity cost of *cx*_*i*_, where *c* is the marginal cost of signal production.

The signal produced by the initially attacked individual is transferred to the other *N*-1 individuals on the patch (receivers). We make no assumptions about how the signal is transferred, so it may be transferred through a common mycorrhizal network, the air, or any other mechanism (17, 34). The signal warns the receivers that an attack is imminent.

Consequently, the receivers can prepare for being attacked, and defend themselves, meaning they suffer a reduced fecundity cost of being attacked. We assume that the timing of herbivore-attack in a given generation is not predictable, which means that individuals cannot prepare themselves for herbivore-attack unless they have received a warning signal (43, 44). The extent to which signal-receivers, *j*, respond to the signal, preparing for herbivore-attack, rather than ignoring the signal, is given by *y*_*j*_. Preparation for being attacked (defence) incurs a fecundity cost of *sx*_*i*_*y*_*j*_, where *s* is the marginal cost of defence, but it reduces the cost of being attacked, which is now given by *d(1-x*_*i*_*y*_*j*_). Note that *x*_*i*_ (signal investment by the signaller) features in these costs to the signal-receiver because, to respond to a signal (mediated by *y*_*j*_), there needs to be a signal there to respond to (mediated by *x*_*i*_), hence why *x*_*i*_ and *y*_*j*_ are multiplied together. We assume that the cost of defence is less than being attacked (*s<d*), which ensures that individuals are favoured to defend themselves against attack (rather than let themselves be damaged).

We then allow individuals to produce offspring (juvenile haploid clones) in proportion to their fecundity. A random sample of *N* juvenile individuals are chosen, for each patch, to survive and form the next adult population (local population regulation). This generates competition between juveniles to obtain a spot on the patch to grow into an adult (Appendix O). After population regulation, a proportion of the new adult population, *m*, migrate to different and random patches. The remaining proportion, *1-m*, remain on their local patch. This lifecycle then iterates over many generations until an evolutionary end point is reached. In Appendix L, we determined the equilibrium (ESS) levels of signalling investment (*x\**) and signal-response (*y\**).

#### Signalling is not evolutionarily stable

We found that individual plants were not favoured to produce warning signals (*x\**=0) (Fig. 2). Any tendency to honestly signal an impending attack will ultimately be removed by natural selection, to avoid providing a benefit to competitors. Honest signalling is a helping behaviour, that increases the fitness of social partners by warning them about an impending attack. Helping can be potentially favoured if it is directed towards relatives that share genes for helping, termed kin selection (28, 45). When the migration rate (*m*) is lower, individuals on a patch will be more closely related. However, a lower migration rate also means that the individuals receiving help are in stronger local competition with the helper, which negates the benefit of helping relatives (29). Put simply, there is no benefit in helping one’s relative if this comes at an equal cost to another relative (30, 46, 47). We showed in Appendices L, M & O that this result also holds if: population regulation occurs after migration, rather than before; plants signal even if they are not the first plant on the patch to be attacked (obligate versus facultative signalling); competition affects fecundity rather than survival.

**Figure 2.**
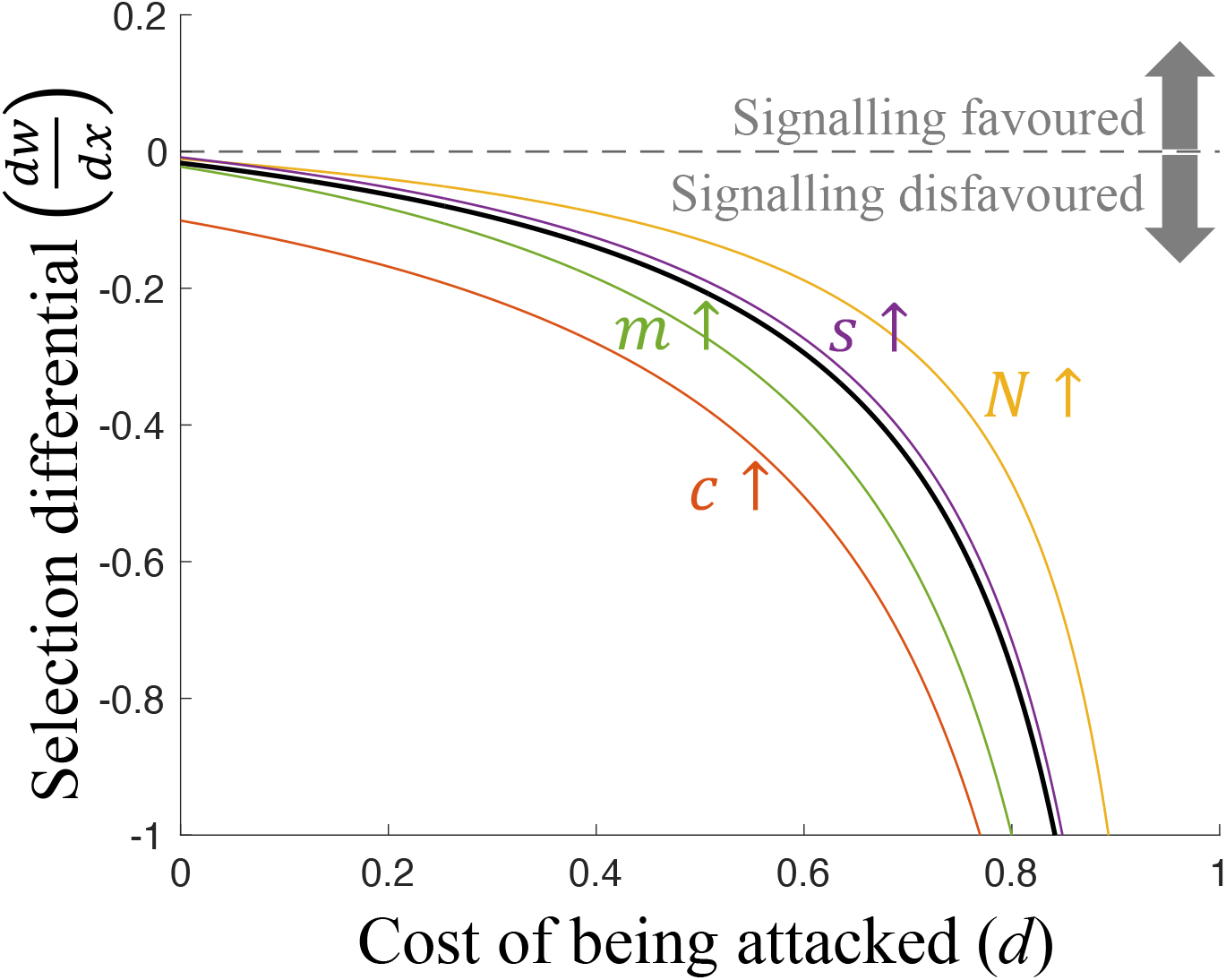
Plant signalling is not evolutionarily stable. Signalling is favoured if plant fitness (*w*) increases with signalling investment (*x*), in other words if the selection differential 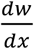 is positive. The solid black line is a reference line that plots the selection differential for a given set of parameter values. The black line never goes above zero, showing that, for our set of parameter values, the selection differential is negative, meaning signalling is disfavoured. Each coloured lines plots the selection differential when one parameter (annotated) is changed to a higher value. The coloured lines do not lie on top of the black line, but like the black line, they never go above zero. This illustrates that changing parameter values can quantitatively adjust the intensity of selection on signalling, but it cannot make the selection differential positive. Signalling is therefore disfavoured for all parameter combinations. We assumed: (*parameter values used for the black line*) *N*=3 (group size), *c*=0.1 (cost of signalling), *s*=0 (defence cost), *m*=0.3 (migration rate); (*alternative parameter values used for the coloured lines*) *N*=7, *g*=1, c=0.6, *s*=0.05, *m*=1; *x*=0, *y*=1, *z*=0 (signals initially rare, honest and responded-to).

#### Alternative population structures

Our model assumed a simple life history where offspring (seeds) can disperse after reproduction. This is reasonable for plants, where the lack of movement after they have started growing means that any neighbours potentially helped will tend also to be competitors (48–53). We also assumed that the parameters of the life cycle such as the migration rate determined both the relatedness (genetic similarity) between different individuals on a patch and who competition occurs between. As an alternative approach, we can construct an ‘open’ model that detaches these model parameters and keeps them as free variables (54). While this can be artificial, it can suggest the kind of conditions that would be required for honest signalling to be favoured. We showed in Appendix M that honest signalling can be favoured when signallers are highly related to signal-receivers but not to their competitors. Although theoretically this is possible, there may not be many scenarios in which low dispersal could lead to high relatedness among interacting plants, but not local competition (48, 49, 51–53). An alternative possibility is if there is a way to preferentially signal to related plants (kin discrimination) (55, 56). We show in Appendix E that kin discrimination can allow honest warning signals to be evolutionarily stable. However, the extent to which plants can discriminate kin, especially with warning signals, is a matter of empirical debate (32, 57, 58). Recent theoretical work has shown how kin discrimination could be favoured if herbivory information is transferred through volatiles (50). Other hypotheses for honest signalling that could be tested empirically include certain forms of generation overlap; or a private (direct) benefit to helping reduce the local population of herbivores (28, 54, 59).

#### Can dishonest signalling be favoured?

Our model also allowed us to investigate the opposite of honest signalling – whether plants could be favoured to produce dishonest signals, where they signalled an attack when this had not happened. This could potentially benefit the dishonestly-signalling plant if it sufficiently reduced the fecundity of their local competitors. In generations where the population is not attacked, a random individual on each deme, *i*, is given the opportunity to signal an attack even though no attack had occurred (dishonest signalling). This individual invests *cx*_*i*_*z*_*i*_ into signal production, where *z*_*i*_ denotes dishonesty. The receivers, *j*, consequently suffer a defence cost of *sx*_*i*_*y*_*j*_*z*_*i*_. In Appendix L, we determined the equilibrium (ESS) level of signal dishonesty (*z*\*) and how it influences signal-response (*y\**).

We found that individuals could be favoured to produce dishonest signals, in which they signalled an attack even when none had occurred. Plants could gain a benefit from dishonest signalling because it harms their local competitors. However, this leads to selection on the receiver plants to ignore signals, and so dishonest signalling would only be transient, not evolutionarily stable. Dishonest signalling has not been empirically observed, but it has also not been tested for. More generally, dishonest signals can in principle be stable within signalling systems when they are at a low frequency, and so do not completely remove the benefit of responding to signals. For example, fork-tailed drongos produce alarm calls to signal to meerkats when a predator is approaching; but also occasionally produce false alarm calls, in the absence of a predator, to steal food left by the fleeing meerkats (60, 61).

### Cue-suppression model

We then examined the case where a plant produces a cue when it is being attacked. This would include chemicals that are produced or released by damage from herbivores or pathogens, and which are transmitted by any route, including the air or fungal network (62–75). The production of a cue could provide another explanation for the experimental data showing upregulation of defences by the neighbours of attacked plants. Can the plant be selected to suppress this cue? (33) We test this by modifying the assumptions of the previous model so that now, after an individual is attacked, it releases a *cue* rather than a *signal*. The cue indicates to the other *N*-1 individuals on the patch (receivers) that an attack is imminent. However, we assume that the attacked plant may pay a cost to suppress the cue, to stop the receivers from learning that an attack is imminent. So now, *x*_*i*_ denotes investment into cue-suppression, rather than signalling. We assume that plants respond to cues by upregulating their defences. In Appendix G, we determined the equilibrium (ESS) level of cue-suppression (*x\**).

#### Cue-suppression is evolutionarily stable

We found that individuals could be favoured to suppress cues (information) of an attack (Fig. 3a). Plants can gain a benefit from cue-suppression because it harms their local competitors (by avoiding helping them). Cue-suppression is favoured when the benefit incurred due to competitors being less able to defend themselves against the attack is smaller than the cost of suppressing the cue (Fig. 3a). More generally, we showed in Appendix H that cue-suppression is favoured with low relatedness between individuals on a patch and local competition. However, since there are many situations in which complete cue suppression is not favoured, the detection of a cue that another plant is being attacked remains an evolutionarily viable explanation for the experimental data (27). For instance, in nature, it may be too costly or even impossible to suppress all cues of herbivore-attack, given the sheer volume and diversity of such cues, passed through the air or fungal networks (34, 76).

**Figure 3.**
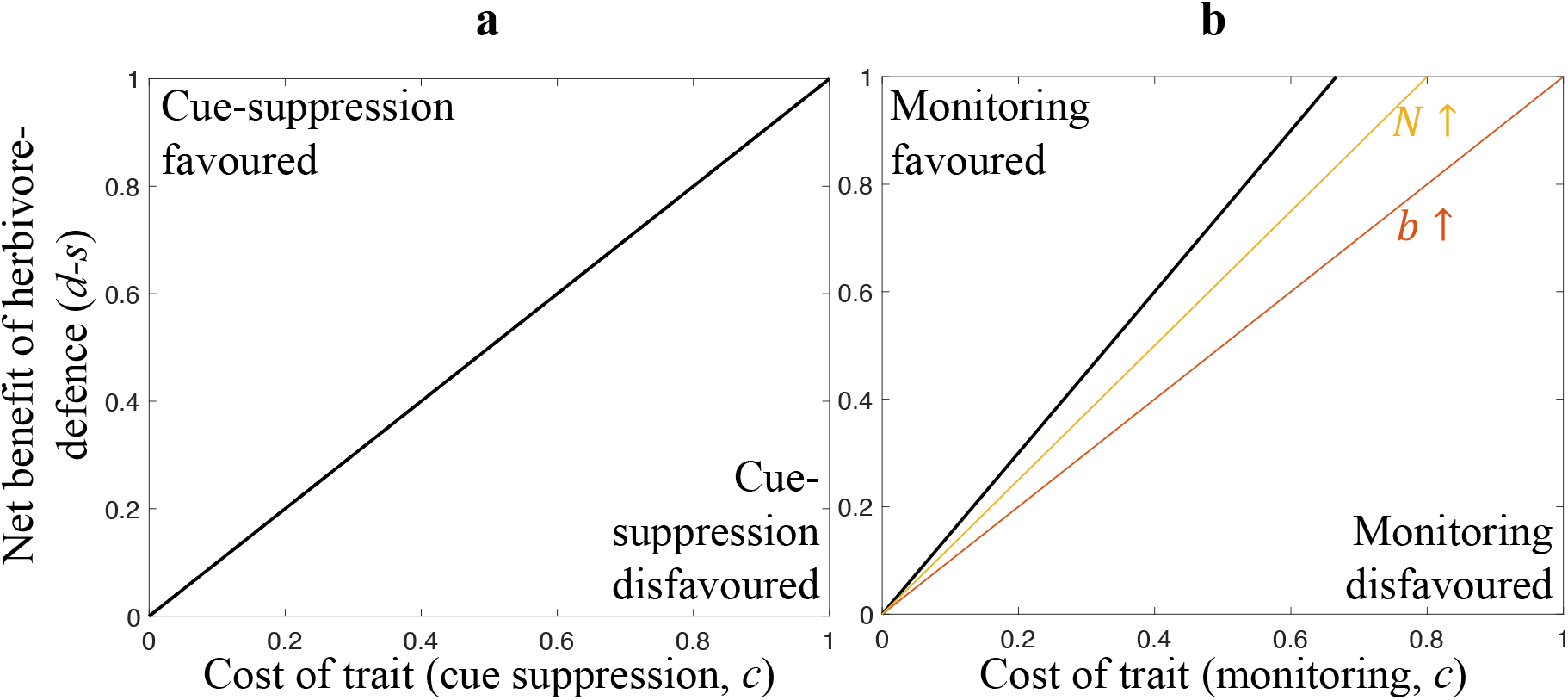
The evolution of cue-suppression and monitoring. Traits are favoured above the lines and disfavoured below them (*i*.*e*. the lines mark the boundary where the trait changes from being favoured / disfavoured, for a given set of parameter values). Solid black lines show boundaries that arise with a given set of parameter values. Coloured lines show boundaries that arise if one parameter is changed to a higher value. A flatter (lower gradient) boundary line implies that the trait is favoured over a larger range of parameter values. Therefore, if a coloured line is flatter than the black line, this implies that an increase in the value of the annotated parameter causes the trait to be favoured more permissively (*i*.*e*., over a greater range of parameter combinations). (**A**) Suppression of cues of herbivore-presence is favoured when the net benefit of herbivore-defence (*d*-*s*) is greater than the trait cost (*c*). (**B**) Mycorrhizal monitoring and signalling is more likely to be favoured when: the benefit of high-quality trade-partners (*b*), *N* (group size) & *d* (attack cost) are high; *s* (defence cost) & *c* are low. We assumed: (A) any value for *N*; (B, *reference values*) *N*=3, *b*=1; (B, *alternative values*) *N*=5, *b*=1.5.

### Fungal monitoring model

Finally, we examine selection on mycorrhizal fungi and ask whether they can be selected to monitor plants, to determine whether they are being attacked, and then signal that information to the other plants they are connected to (13). To examine this, we modify the assumptions of the first ‘plant signalling’ model, so that now, as well as the (infinite) population of plants, we additionally model an infinite population of fungi.

The population is split into patches comprising *N* plants connected by one fungus (which means that each plant is associated with just one fungus). When a plant is attacked, the fungus may monitor cues of herbivore attack produced by the initially attacked plant and signal this information to the other plants it is connected to (recipients). As before, signals may be dishonest and ignored. The difference is that, now, signal investment and signal dishonesty are fungal rather than plant traits (signal response is still a plant trait). We assume that fungi gain a benefit by being connected to plants in better condition (‘fitter’), because such plants will be more able to provide the fungus with carbon in exchange for nutrients provided by the fungus (i.e. these plants are ‘better trade partners’) (35, 36, 77). Specifically, a fungus gains a benefit of *bf*_*partner*_, where *f*_*partner*_ gives the average condition (fecundity) of the plants it is connected to. Fungi reproduce by producing offspring (juveniles) in proportion to their fecundity; each juvenile migrates to a random deme, and a random juvenile is chosen for each deme to survive and form the next adult fungal population. In Appendix N, we determined the equilibrium (ESS) levels of fungal signaling investment 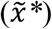 and plant signal-response (*y\**).

#### Fungal monitoring & signalling is evolutionarily stable

We found that mycorrhizal fungi could be selected to monitor plants, to determine whether they are being attacked, and then signal that information to (warn) the other plants they are connected to. We found that fungi could gain a benefit from this, because it helps their trade partners defend themselves against attacks, allowing them to better transfer carbon to the fungus. Mathematically, fungal monitoring and signalling is favoured when the costs to the fungus of monitoring and signalling are less than the benefit to the fungus of having higher quality trade partners (Fig. 3b). In contrast, we showed in Appendix N that fungi are never favoured to signal dishonestly, by tricking plants into upregulating their defences when no attack is imminent, because this would reduce the quality of their trade partners. Empirically, mycorrhizal fungi are capable of both perceiving cues of herbivore-attack and inducing herbivore resistance in the plants they are connected to (16, 78).

One possibility we have not considered is if multiple fungal individuals associate with each other or the same plant (79, 80). Previous theory has shown that when more fungi and plants interact within a network this favours more efficient resource trading, and hence helps make trading evolutionarily stable (77). More complex network interactions could similarly influence the evolution of monitoring and signalling by fungi. For example, could larger networks lead to fungi monitoring other fungi, and hence increased selection for fungal signalling? This would be especially the case if it helped facilitate trading across the network. Or could larger networks select for fungal cheating, where some fungi avoid any cost of signalling? Answering such questions would require an alternative modelling approach, that examined more complex networks. Nonetheless, more complex situations do not change that fungi can gain a benefit from monitoring plants and then passing that information to other plants that they trade with.

## Discussion

We have applied a body of signalling theory that has been well-developed to explore animal behaviour to a new context, information transfer between plants. Our results show that plants: (1) are unlikely to be selected to signal to their neighbours about the presence of herbivores (Fig. 2); (2) can be favoured to produce dishonest signals, where they signal an attack when none has occurred, but that this will select for other plants to ignore signals; (3) can be favoured to pay resources to suppress any information (cue) to neighbours about attack (Fig. 3a). In contrast, mycorrhizal fungi can be selected to monitor their host plants, detect when they are attacked, and then signal (warn) other plants in their network (Fig. 3b).

Our results do not support the hypothesis of warning signals by plants, passed *via* any route, including common mycorrhizal networks or the air (2, 11–24, 27, 81). This is because these warnings would benefit neighbouring competitors, to the cost of the signalling individual. Furthermore, we found that not only are plants not expected to signal, but that they can be selected to signal dishonestly or to actively suppress any cues of being attacked. Dishonest signalling or suppression of cues is favoured to harm or avoid helping neighbouring competitors. Empirically, there is little evidence for plant–plant honest signalling (intraspecific), though plants can be favoured to signal honestly to their pollinators and seed dispersers (interspecific), who they are not in direct competition with (34, 82–86).

For a helping behaviour to be favoured, such as signalling a warning of herbivore or pathogen attack, we showed that this would require that helping and competition occur at different scales (economic neighbourhoods) or some method of kin recognition / discrimination (30, 31, 46, 47, 55, 87–89). Helping and competition occurring at different scales is relatively unlikely for plants because they are immobile, meaning local interactions involve both cooperation and competition for resources, disfavouring warning signalling (90). This is not always the case in mobile organisms such as animals and bacteria. The same problem of local competition has been demonstrated empirically in other organisms, such as when fig wasps compete for mates in the closed environment of a fig fruit, or when bacteria secrete ‘public goods’ (91–93). However, the problem of local competition can be overcome in mobile organisms, if helping occurs between relatives before they disperse to compete with non-relatives (94–96). In animals, warnings about the presence of predators have also been argued to be favoured because they also reduce predation on the individual making the warning call – this is different from plants, where the warning arises after attack (97–99). In contrast, local competition favours harming behaviours, because harming neighbours can decrease competition for resources (31, 100–102). Alternative modelling approaches to examine these issues could include explicit spatial structures, such as on a graph or lattice, but these have been shown to lead to analogous results (87, 103–105).

Information about herbivore-attack could potentially occur through mycorrhizal networks or volatiles in the air (33, 34, 74). The production of herbivore-induced plant volatiles may be unavoidable and does not seem to confer a fitness benefit on the producer (34), leading to the suggestion that herbivore-induced plant volatiles are likely to represent cues rather than signals (62–75). This is consistent with our plant signalling and cue-suppression models, which did not make any assumptions about how information about herbivore-attack is transferred. More generally, there may be many ways for information about herbivore-attack to be transferred, suppressed, directed towards kin, *etc*., with some mechanisms more biologically plausible than others (34). Our intention has been to examine the evolutionary stability of different forms of information transfer, in a way that could be applied to a diversity of proximate mechanisms (106).

In contrast to the situation for plants, we found that mycorrhizal fungi can be favoured to monitor their host plants, detect a cue of when they are attacked, and then signal this to (warn) other plants in their network (13). Fungi are selected to monitor and signal because defended plants will maintain better condition and hence become better trade partners. Previous theory has shown that selection for fungi and plants to trade resources with each other is increased when multiple plants and fungi interact in the same network, because this stabilises efficient trading (77).

To conclude, we examined hypotheses explaining the empirical result that when one plant in a mycorrhizal network is attacked that this leads to other plants in the network upregulating their defence mechanisms (1–5, 8). Our modelling suggests that this is more likely to represent either: a cue produced by plants that is too costly to suppress, or fungi monitoring plants, and then signalling to other plants. Further experiments could test between these possibilities, by examining the underlying mechanism in networks or experimental multiple root systems. How is information conveyed? Where does that information arise from? What are the fitness consequences for all the individuals involved? A greater understanding of these mechanisms could potentially also be exploited in an agricultural context, by facilitating plant defence against herbivores.

## Materials and methods

In the Supplementary Information, comprising Appendices A–O, we analyse and interpret a series of models. The models differ from each other in the assumptions they make about: lifecycle; how demography affects relatedness; what traits can evolve. Appendices L & M present our most general plant signalling models, where signals can evolve to be dishonest and / or ignored. Appendices A–D present special cases of these models, where signals are forced to be honest and responded-to. Appendix E presents a model of plant signalling in which plants can recognise their kin. Appendices G–J present cue-suppression models. Appendix N presents fungal monitoring models. Appendix F provides some illustrative ‘inclusive fitness’ versions of our models. Appendix K interprets the models presented in Appendices A–J, setting them in the context of the wider literature on the evolution of helping and harming. Appendix O provides some supplementary discussion of how ‘competition’ is modelled.

## Acknowledgements

We thank Anna Dewar, Ashleigh Griffin, Jason Hoeksema, Vasilis Kokkoris, Asher Leeks, George Shillcock and two anonymous reviewers for comments on the manuscript. We thank the ERC (834164 & 771387), HFSP Program Grant (RGP 0029), NWO-VICI (202.012), NWO-Microp program, Hefner Foundation, and Ammodo Foundation for funding. Figure 1 was made with https://Biorender.com.

## Supplementary Information

### Overview

In this Supplementary Information, we analyse and interpret a series of models. The main signalling model presented in the main text is analysed in Appendix L. Readers interested in the details of this model should skip straight to this appendix. A variation of this model (with an alternative lifecycle assumption) is analysed in Appendix M.

Simplified versions of the signalling models are analysed in Appendices A–D. These models are not as general as the full signalling model presented in the main text and in Appendices L & M. Specifically, in these simplified models, signals are forced to be honest, and individuals are forced to respond to signals (both of these assumptions are relaxed in the full models). We include these simplified models for two reasons. Firstly, their mathematical simplicity may help some readers get a sense of how the models are working, before tackling the full models. Secondly, they allow us to make clear links to the wider social evolution literature on the evolution of helping and harming; we make these links in Appendix K.

The ‘kin discrimination’ signalling model is analysed in Appendix E. The main cue-suppression model referred to in the main text is analysed in Appendix G (variations of this model are analysed in Appendices H, I & J). The fungal monitoring model referred to in the main text is analysed in Appendix N.

Most of our models are formulated using ‘neighbour-modulated’ fitness rather than ‘inclusive’ fitness (1). The two fitness measures are formally equivalent under a standard set of mathematical assumptions, and so neighbour-modulated models are usually interpreted from an inclusive fitness perspective (2–4). However, inclusive fitness analyses can be more intuitive (5), and so we have provided some illustrative inclusive fitness analyses in Appendix F, to complement our main neighbour-modulated fitness approach.

We provide some supplementary discussion of how ‘competition’ is modelled in Appendix O.

### Open and closed models

Some of our models assume (accurately) that relatedness emerges from model parameters (closed models), and others artificially ‘detach’ relatedness from lifecycle assumptions so that it can be treated as a parameter (open models) (4, 6). The reason why we analysed (technically inaccurate) open models as well as (accurate) closed models is that, in closed models, ‘who you socially interact with’ and ‘who you compete with’ are both determined by model parameters, meaning they are correlated. This makes the effects of kin selection (favouring helping behaviours) and kin competition (disfavouring helping behaviours) hard to disentangle (4, 5, 7–11). Open models detach relatedness from model parameters, so that the causal impact of kin selection can be more easily seen, free from associated competitive effects. Examining open models alongside closed ones has a rich tradition in social evolution research (4, 6).

### Alternative lifecycle assumptions

Some of our models assume a lifecycle where population regulation occurs before migration, and others assume that population regulation occurs after migration (7). The reason why we analysed ‘population regulation before migration’ and ‘population regulation after migration’ versions of the models is that, in the former case, individuals only compete with individuals on their native patch, but in the latter case, individuals compete to some extent with individuals on other patches. This is known to have potentially dramatic consequences on the type of social behaviour that evolves (7). It is therefore important to examine how sensitive conclusions are to lifecycle assumptions.

### Data Accessibility

A *Mathematica* file comprising all calculations in this Supplementary Information, and a *Matlab* file implementing the numerical procedure used to solve Model M (Appendix M), can be accessed at https://github.com/ThomasWilliamScott/Plant_Signalling. The results figures in the main text plot results for Model L (signalling; Fig. 2), Model G (cue-suppression; Fig. 3a), and Model N (monitoring; Fig. 3b).

## Appendix A

### Model A: signalling; closed; population regulation before migration

We assume an infinite population of individuals (*e*.*g*., plants). We assume for simplicity that individuals are haploid and asexual. The population is split into demes of size *N* (infinite island model) (12). We focus our analysis on a focal individual, drawn at random from the population.

Each generation, a random individual on each deme is initially attacked by a herbivore, and suffers a fecundity cost of *d*. Given that there are *N* individuals on each deme, each individual has a 1/*N* chance of being initially attacked by the herbivore. The initially-attacked individual may then invest into the production of a signal. If the focal individual is the initially-attacked individual on its deme, it invests an amount *x*_*focal*_ into signal production. Signal investment incurs a fecundity cost that scales linearly with *c*. Therefore, the focal individual pays a signalling cost of *cx*_*focal*_ when it is initially-attacked.

The signal produced by the initially-attacked individual is transferred to (and recognised by) the other *N*-1 individuals on the deme. We make no assumptions about how the signal is transferred, but it may be being transferred through a common mycorrhizal network, or through the air, *etc*. The signal warns the other plants that a herbivore is in the vicinity. Consequently, the other plants can prepare for being attacked, and defend themselves, meaning they suffer a reduced fecundity cost of attack. Preparation for herbivore attack (defence) incurs a fecundity cost that scales linearly with *s*. We denote the average signalling investment amongst the *N*-1 other plants on the focal individual’s deme by *x*_*others*_. This means that, when the focal individual is not the initially-attacked individual, it will ultimately suffer a cost of attack that is given by *d(1-x*_*others*_), and a defence cost that is given by *sx*_*others*_. We assume that preparing for herbivore attack is less costly than the fecundity cost of being attacked (*d*>*s*), which ensures that individuals are favoured to defend themselves against herbivores.

We emphasise that, in our model, only the initially-attacked individual has a chance to warn the other individuals about the presence of a herbivore. Subsequently-attacked individuals may still produce the signal, but our assumption is that by then, the herbivore has already arrived, rendering the signal useless for warning others about herbivore presence. We allow for the possibility that subsequently-attacked (*i*.*e*., not initially-attacked) individuals can facultatively turn off signal production, so that they do not pay the costs of producing a useless signal. We assume that the focal individual pays a signalling cost of *cx*_*focal*_*(1-g)* when subsequently-attacked, where *g* denotes the extent to which signal investment is ‘facultative’ as opposed to ‘obligate’ (analogously, a non-focal individual pays a signalling cost of *cx*_*others*_*(1-g)* when subsequently-attacked). A fully obligate signal (*g*=0) means that it is produced irrespective of whether the individual is initially-attacked (rendering the signal is useful) or subsequently-attacked (rendering the signal useless).

After the fecundity costs of attack and signal investment have been incurred by all individuals, each individual produces a large number of offspring (juvenile haploid clones) in proportion to their fecundity. A random sample of *N* juvenile individuals are chosen, for each deme, to survive and form the next adult population (local population regulation). After population regulation, a proportion of the (new) adult population, *m*, migrate to a different deme, randomly chosen for each individual. The remaining proportion, *1-m*, remain on their local deme. This lifecycle then iterates over many generations until an evolutionary end point is reached.

We denote the population average investment into signalling by *x*_*pop*_. We denote the average signalling investment on the focal individual’s deme (incorporating its own signalling investment) by *x*_*deme*_; this can be written in terms of *x*_*focal*_ and *x*_*others*_ as

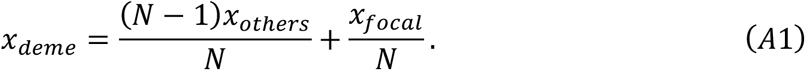

We assume that the baseline fecundity is 1, that 0 ≤ *x*_*focal*_, *x*_*others*_, *x*_*deme*_, *x*_*pop*_ ≤ 1, and that *c* + *d* + *s* ≤ 1; these assumptions ensure that fecundity never falls below zero. The number of juvenile offspring produced by the focal individual is then proportional to

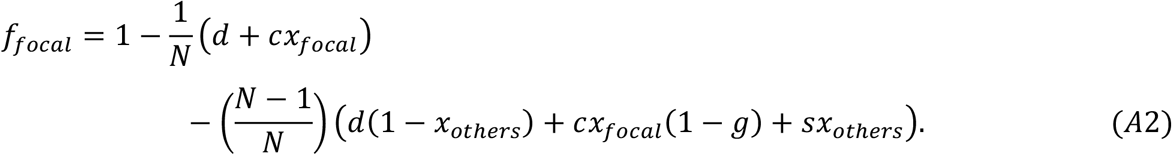

We can make sense of the right-hand side of Equation A2 as follows. Baseline fecundity is one (left-hand term). With probability 1/*N*, the focal individual is attacked first, leading to additively applied costs of attack (*d*) and signal production (*cx*_*focal*_), leading to the second term. With probability 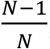, the focal individual is not attacked first, and can receive a signal of herbivore presence from another individual on the deme, leading to additively applied costs of attack (*d*(*1-x*_*others*_)) and defence (*sx*_*others*_), plus the costs of signal production that are incurred even when the focal individual is not attacked first, owing to obligatory (unconditional) expression (*cx*_*focal*_(*1-g*)). This leads to the third term.

The number of juvenile offspring produced by an individual drawn randomly from the focal individual’s deme (including the focal individual) is proportional to

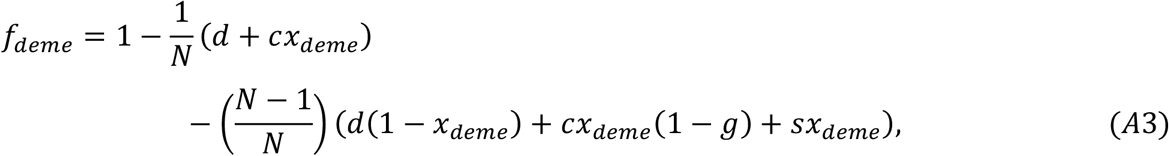

which we can rewrite using Equation A1 (getting rid of the superfluous *x*_*deme*_ parameter) as

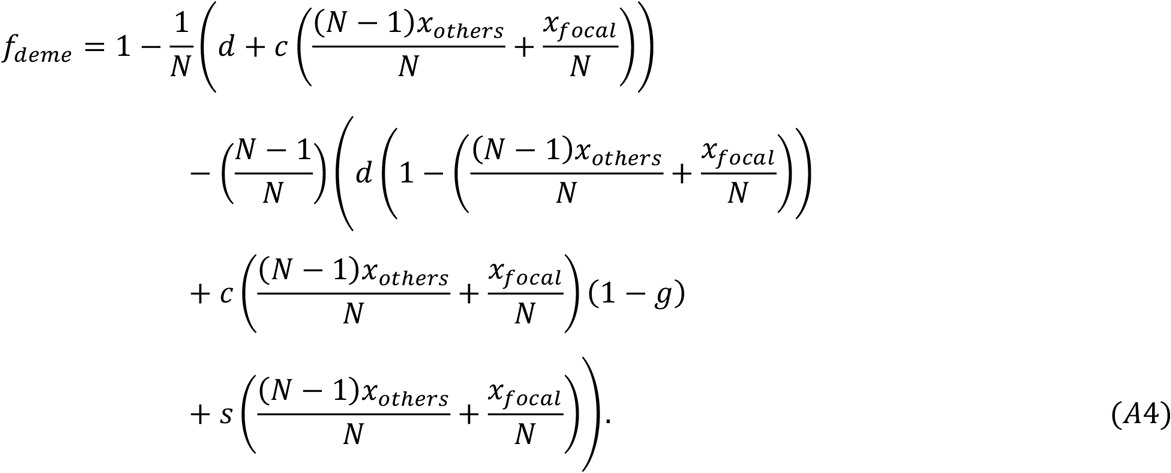

The number of juvenile offspring produced by an individual drawn randomly from the population is proportional to

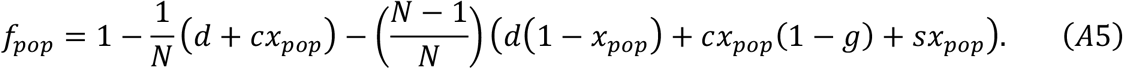

The fitness of the focal individual is then given by:

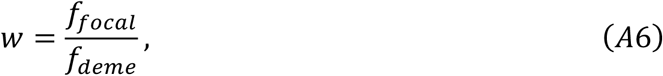

where explicit expressions for the fecundity (*f*_*focal*_, *f*_*deme*_) functions are provided in Equations A2 & A4 (7, 12, 13). Note that the *f*_*pop*_ function derived above (Equation A5) does not feature in this fitness function, but it will feature in the fitness function of some subsequent models. ‘Fitness’ refers to the number of surviving offspring produced by an individual after one full iteration of the lifecycle (12). Under our lifecycle assumptions, the size of the population remains constant after each full iteration of the lifecycle, and this is reflected by the fact that the average fitness taken over the population is equal to 1.

As a technical aside, we note that, by ‘fitness’, we strictly mean ‘neighbour-modulated’ fitness, which counts up fitness effects on a focal individual. This can be conceptually distinguished from ‘inclusive’ fitness, which counts up fitness effects arising from the actions of a focal individual, though the two fitness measures are formally equivalent under a standard set of mathematical assumptions (2, 12, 14–20). We have provided some illustrative inclusive fitness analyses in Appendix F.

Equation A6 reflects the fact that, under our assumption that (local) population regulation occurs before migration, individuals only compete with competitors on their native deme, rather than with individuals on other demes. This is why the fecundity of the focal individual (*f*_*focal*_) is weighted only by the average fecundity of an individual from the native deme (*f*_*deme*_), rather than by the average fecundity of an individual drawn from the population at large (*f*_*pop*_).

Our aim is to examine what level of signal investment evolves at evolutionary equilibrium. To do this, we assume a monomorphic population, where all individuals invest the same amount into signal production, given by *x\** (the lack of subscript implies that the value is the same across the population). We then take a random (focal) individual and mutate its (and its identical-by-descent relatives’) level of signal investment to a deviant value *x*, and ask how the fitness of the focal individual changes in response, in effect calculating 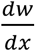. Evaluating 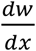 at equilibrium (*x\**), the equilibrium (ESS) level of signal production can then be obtained as: (*i*)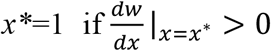 (maximal signal production); (*ii*)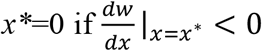 (no signal production); (*iii*) 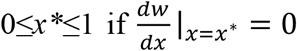, where *x\** is specifically obtained as the value for which 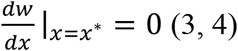.

To calculate 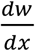, we first expand it using the chain rule

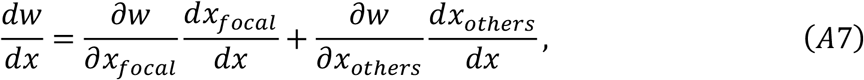

and then proceed to evaluate the derivatives on the right-hand side (3, 4). Because the population is monomorphic, and because we are interested in the long term evolutionary state of the population, we evaluate these derivatives at the point where *x*_*focal*_, *x*_*others*_, *x*_*pop*_ = *x*^*^. We trivially obtain

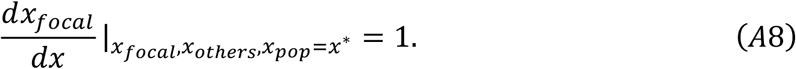

We obtain the equilibrium-evaluated partial derivatives 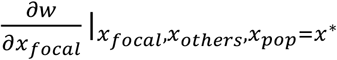 and 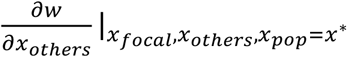respectively as

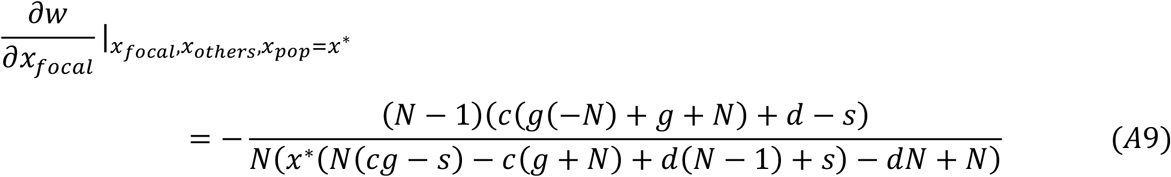

and

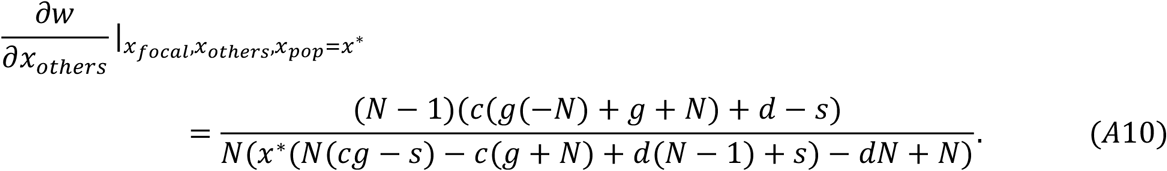

We obtain 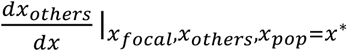 by first noting that 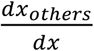 is interpretable as a coefficient of relatedness, since it denotes the marginal (correlated) change in signal production by a social partner as a consequence of a change in signal production by an actor (3, 4). We can therefore write

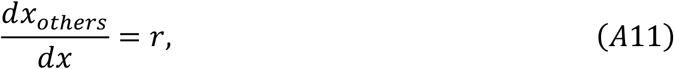

where *r* denotes the relatedness between social partners. We note that *r* is an ‘others-only’ measure of relatedness, as opposed to a ‘whole-group’ measure of relatedness, since it measures the relatedness between an individual and its social partners (who are the other members of the deme), rather than the relatedness between an individual and a random member of its deme (including itself) (21).

We cannot evaluate *r* in general, because it is changing over time and will depend on initial genotype frequencies, but fortunately, we can evaluate it at equilibrium, which is the point that we are interested in anyway, to find

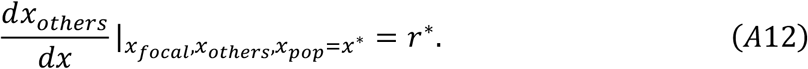

To get an expression for the equilibrium relatedness, *r\**, it is useful to first calculate the equilibrium ‘whole-group’ relatedness for this model, 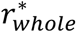, then convert back to find the equilibrium ‘others-only’ relatedness, *r\**. The equilibrium whole-group relatedness can be found by first writing a recursion for how whole group relatedness, *r*_*whole*_, changes across a generation (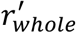 denotes whole-group relatedness in the next generation):

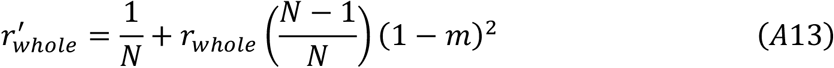

(7, 10, 12). We obtained Equation A13 by calculating the average relatedness between a focal individual and a randomly drawn member of the focal individual’s deme after one iteration of the lifecycle. With probability 1/*N*, the randomly drawn individual is the focal individual itself, leading to a relatedness of 1, hence the first term of the right-hand side of Equation A13. With probability 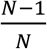, the randomly drawn individual is not the focal individual. With probability 1-*m*, the randomly drawn individual is a non-dispersing individual. With probability 1-*m*, the focal individual is a non-dispersing individual. This leads to a relatedness of *r*_*whole*_, and combining these factors leads to the second term of the right-hand side of Equation A13. With the remining probability 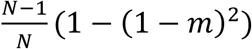, the randomly drawn individual is related to the focal individual by zero, because one of the two individuals have dispersed – the multiplication by zero is why this term does not feature in Equation A13.

Equilibrium whole-group relatedness is then obtained by setting 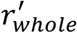 = *r*_*whole*_= 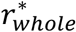 to obtain

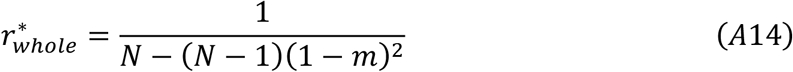

(10). Then, we can write down how whole-group relatedness relates to others-only relatedness:

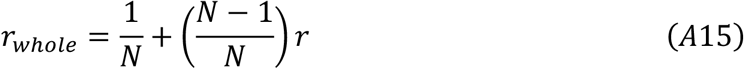

(21). Using Equations A14 & A15, and setting 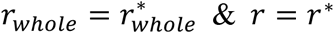, we obtain the equilibrium (others-only) relatedness as

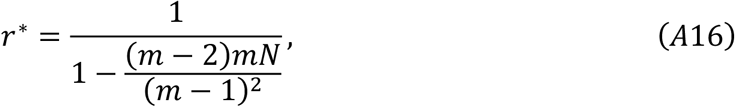

which we can substitute into Equation A12 to find the equilibrium (long-term) value of 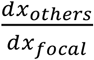:

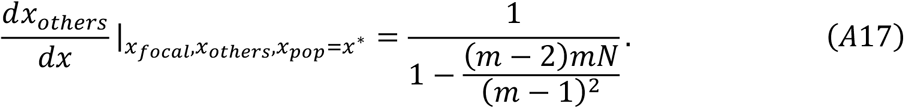

We can now substitute Equations A8, A9, A10 & A17 into Equation A7 to obtain an explicit expression for 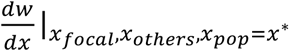:

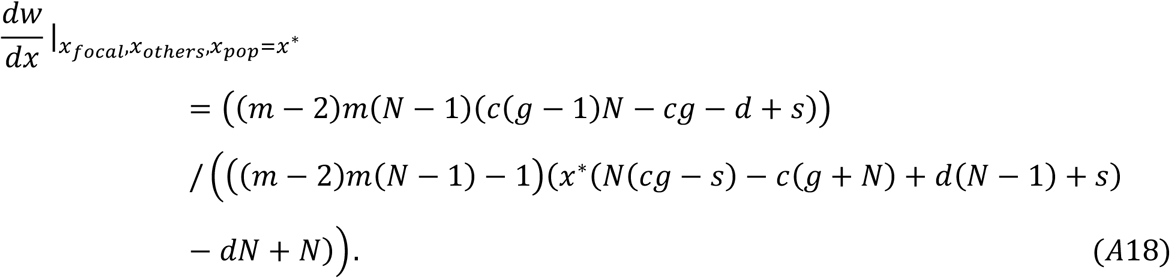

As long as there is some migration (*m*>0), the right-hand side of Equation A18 is strictly negative. If the reader would like to check this for themselves by inspection, remember that most variables are bound between 0 and 1, except for *N* which is greater than or equal to 2, and also remember that *c* + *d* + *s* ≤ 1 and *d* > *s*.

Therefore, in this model, the long-term (ESS) level of signal production is *x*^*^ = 0. At equilibrium, individuals evolve to *not* signal the presence of herbivores to their social partners. Any tendency to signal the presence of herbivores will ultimately be removed by natural selection.

## Appendix B

### Model B: signalling; open; population regulation before migration

We now ‘open’ Model A, detaching relatedness from demographic parameters (*m* and *N*), so that it is now a parameter (constant) in its own right (4, 6). Inspecting Equation A16 reveals that this assumption, of independence of relatedness from demography, is obviously false. But we proceed anyway with this noted.

To analyse this open model, where relatedness varies independently of model parameters, we calculate we calculate 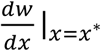 as before, by evaluating the right-hand derivatives of Equation A7 at the point where *x*_*focal*_, *x*_*others*_, *x*_*pop*_ = *x*^*^. The only difference is that, in the previous model, we evaluated the 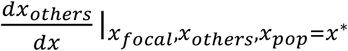 derivative, which is interpretable as an equilibrium value of relatedness, in terms of model parameters *m* and *N*, to obtain Equation A17. However, now, we simply evaluate 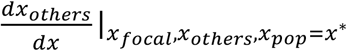 as a constant, *r\**. In other words, in the previous model, we substituted Equation A17 (alongside Equations A8, A9 & A10) into Equation A7 to obtain an explicit expression for 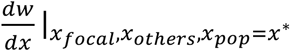, but now, we substitute in Equation A12 instead (still alongside Equations A8, A9 & A10). We obtain

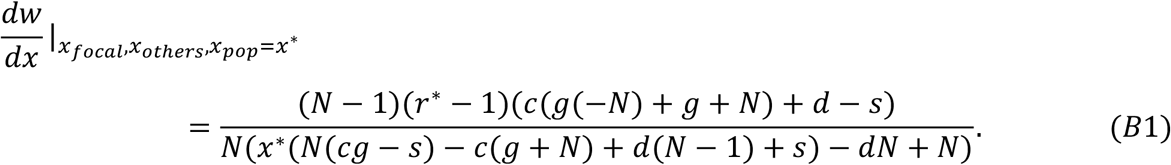

As long as relatedness is not maximal (*r*\*<1), the right-hand side of Equation B1 is strictly negative. Therefore, in this model, the long-term (ESS) level of signal production is *x*^*^ = 0. This is the same result as the equivalent ‘closed’ model (Model A). We have shown that ‘opening’ Model A does not facilitate the evolution of signalling. In the next section, we therefore move onto a slightly different lifecycle, where population regulation occurs after migration, to see if it is more favourable for the evolution of signalling herbivore-presence.

## Appendix C

### Model C: signalling; closed; population regulation after migration

In this Appendix, we formulate and analyse a slightly different version of Model A, where population regulation now occurs after migration, rather than before. Specifically, after each individual has produced a large number of offspring (juvenile haploid clones) in proportion to their fecundity, a proportion of these offspring, *m*, migrate to a different deme, randomly chosen for each juvenile individual. A proportion, *1-m*, remain on the local deme. A random sample of *N* juvenile individuals are then chosen, for each deme, to survive and form the adult population for the next generation (local population regulation).

Under these revised lifecycle assumptions, the fitness of the focal individual is given by

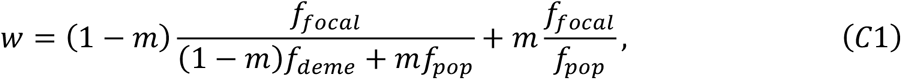

where explicit expressions for the fecundity (*f*_*i*_) functions are provided in Equations A2, A4 & A5 (7, 12, 13).

We can make sense of the first term of the right-hand side of Equation C1 as follows. Of the *f*_*focal*,_ juvenile offspring produced by the focal individual, (1 − *m*)*f*_*focal*,_ of these remain on the native deme, leading to the numerator. These juvenile offspring compete with the (1 − *m*)*f*_*deme*_ non-dispersing juvenile offspring produced by an average member of the native deme, as well as the *mf*_*pop*_ dispersing juvenile offspring produced by an average member of another deme, leading to the denominator. The second term of the right-hand side of Equation C1 can be understood analogously.

We then analyse the model, as before, by evaluating the right-hand derivatives of the 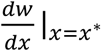 expression (Equation A7) at equilibrium. For this version of the model, these right-hand derivates are calculated as:

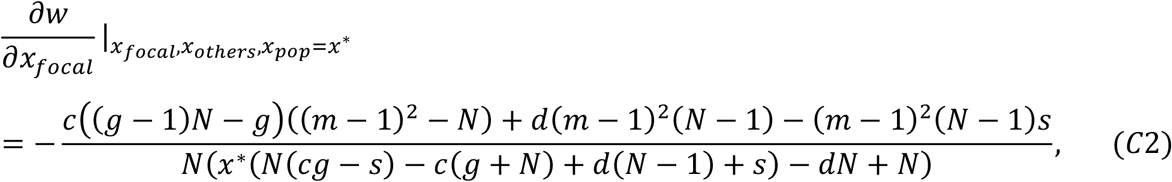

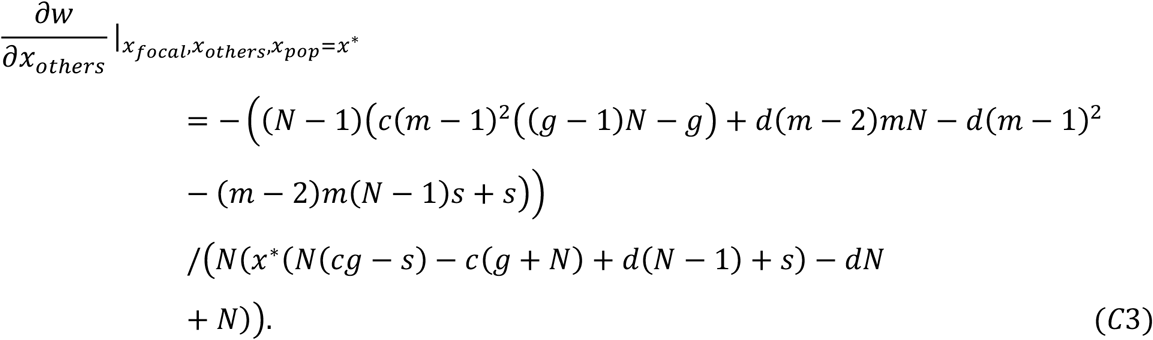

The 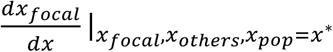 and 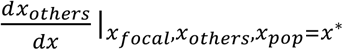 derivatives are still given by Equations A8 and A17 respectively.

By substituting Equations A8, C2, C3 & A17 into Equation A7, we obtain

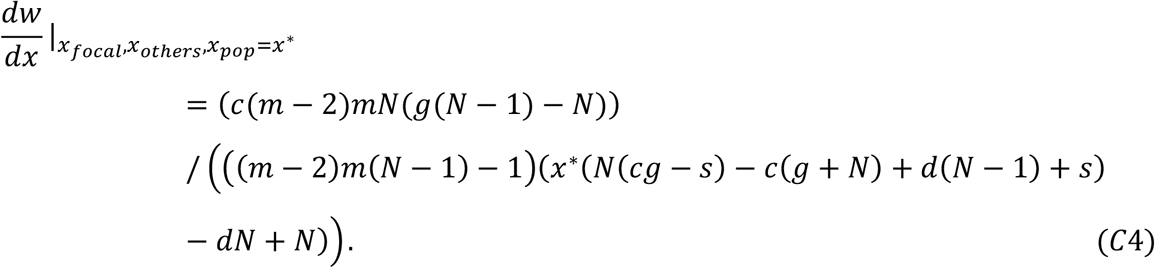

As long as there is some migration (*m*>0) and signalling cost (*c*>0), the right-hand side of Equation C4 is strictly negative. This means that the long-term (ESS) level of signal production is *x*^*^ = 0. This is the same result as Models A & B. Therefore, the lifecycle change (moving population regulation to go after migration) did not facilitate the evolution of signalling. Next, we ‘open’ this model, to see if the lifecycle change is more favourable to the evolution of signalling in an open-model setting.

## Appendix D

### Model D: signalling; open; population regulation after migration

In this Appendix, we formulate and analyse the ‘open’ version of Model C, where relatedness is now treated (artificially) as a constant (4, 6).

As usual, we calculate 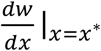 by evaluating its right-hand derivatives (Equation A7). Substituting Equations A8, C2, C3 & A12 into Equation A7, we obtain

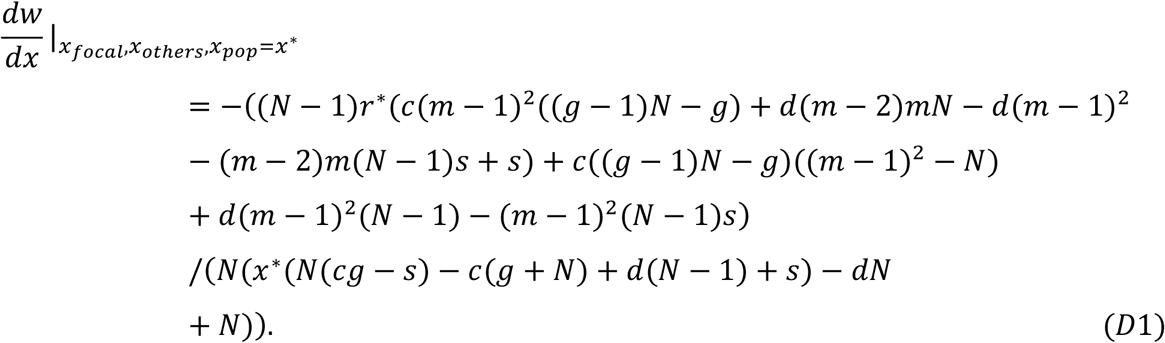

This expression is complex, so we focus on two special cases. First, we consider the case where the migration rate is zero. Substituting *m*=0 into Equation D1, we obtain

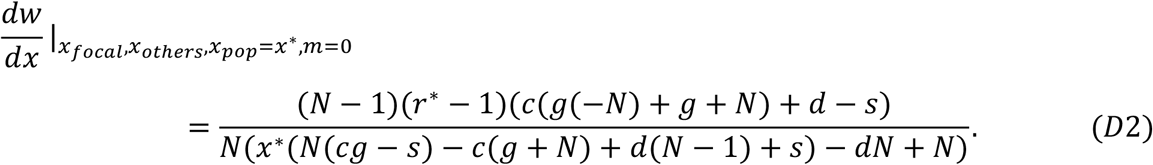

As long as relatedness is not maximal (*r*^*^ < 1), the right-hand side of Equation D2 is strictly negative. This means that, when there is no migration, the long-term (ESS) level of signal production is *x*^*^ = 0 (no signal production).

Second, we consider the case where the migration rate is one. Substituting *m*=1 into Equation D1, we obtain

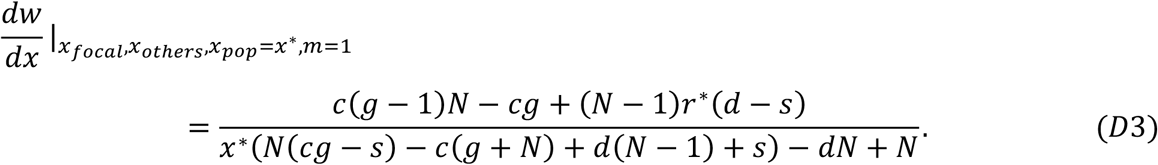

We find that this condition is positive, meaning signalling is favoured (leading to *x\**=1), whenever

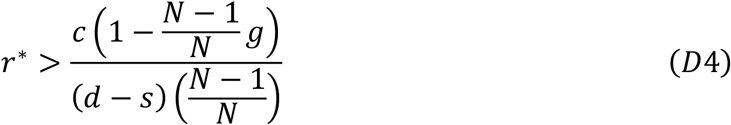

is satisfied. Conversely, signalling is disfavoured, leading to *x\**=0, whenever 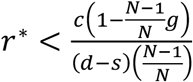 Therefore, signal production is sometimes favoured if there is sufficient migration (*m*).

Equation D4 is a form of Hamilton’s rule, as can be seen by setting 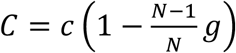 and 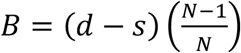 to obtain 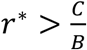. The capital ‘C’ gives the overall cost of the social act (signalling). It is the unit cost of signal production (*c*), weighted by the proportion of incidences where this cost is actually paid 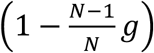. The capital ‘B’ gives the overall benefit of the social act (signalling). It is the unit benefit of signal production, as manifested as a reduced fecundity cost of being attacked (*d*) minus the cost of herbivore defence (*s*), weighted by the proportion of incidences where this benefit is actually received 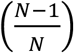.

Therefore, unlike in Models A-C, signalling herbivore-presence can be favoured in this model (if there is sufficient migration and Hamilton’s Rule is satisfied). The combination of ‘population regulation after migration’ and ‘relatedness as a parameter’ (open model) can facilitate the evolution of signalling herbivore-presence.

To give even more intuition for this result, we plot it below in Fig. S1. Signalling is favoured above the lines (technically, this corresponds to when 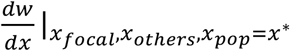, given in Equation D1, is greater than zero).

**Figure S1.**
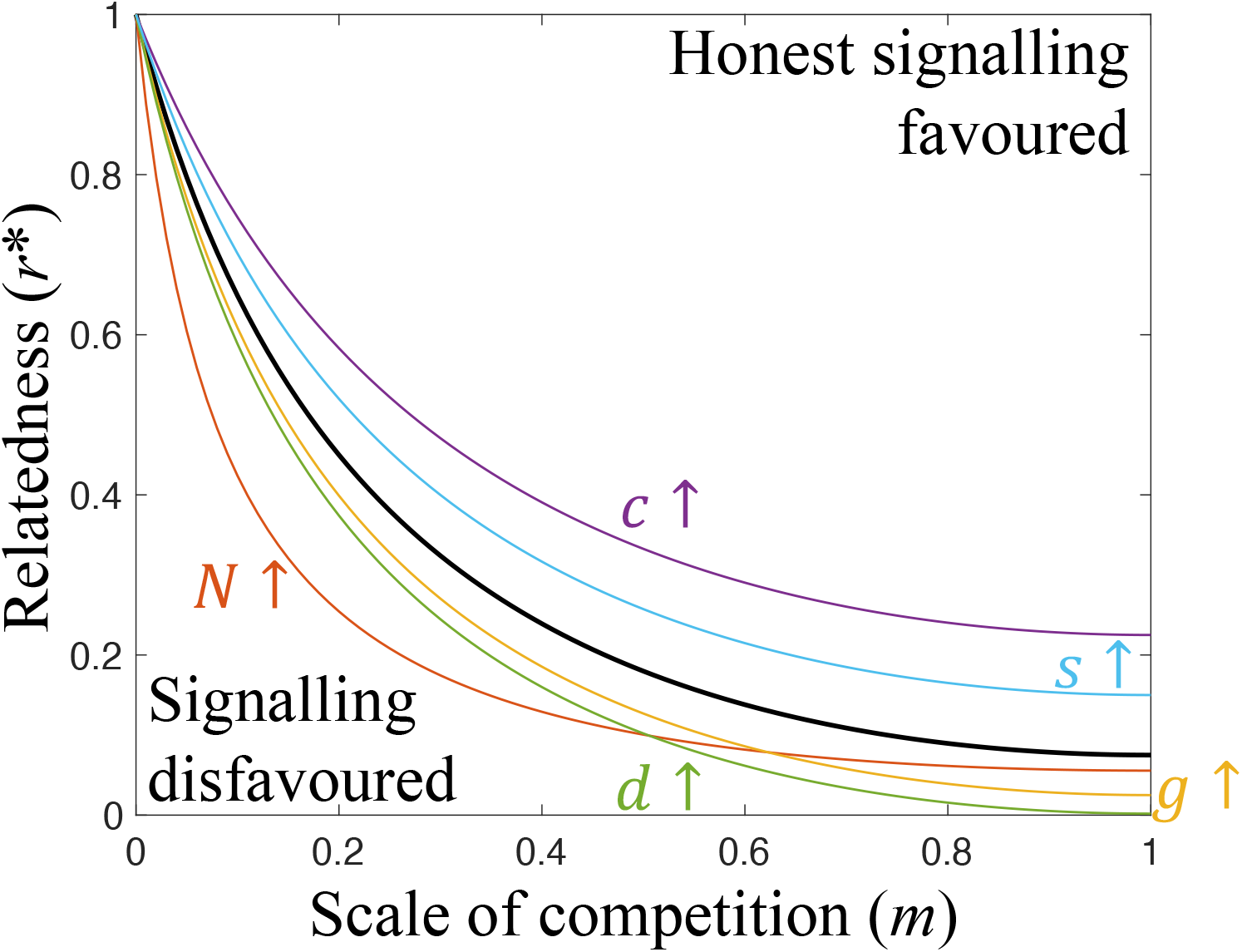
The evolution of signalling in an open model (Model D). Signalling is favoured above the lines and disfavoured below them. Technically, the lines correspond to when 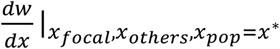 (given in Equation D1) is equal to zero. The solid black line is a reference line generated with arbitrary parameter values. Coloured lines show the effect of increasing parameter values. Signalling is more likely to be favoured when: relatedness (*r*\*), scale of competition (*m*), group size (*N*), attack cost (*d*) and conditionality (*g*) are high; trait cost (*c*) and defence cost (*s*) are low. We assumed: (*reference values*) *d*=0.1, *c*=0.001, *N*=3, *s*=0.08, *g*=0; (*alternative values*) *d*=0.9, *c*=0.003, *N*=10, *s*=0.09, *g*=1.

## Appendix E

### Model E: signalling; closed with kin discrimination; population regulation after migration

In Appendix D (Model D), we found that signalling can evolve if population regulation occurs after migration and if relatedness between social partners is high enough relative to other model parameters (Equation D4). This was an ‘open’ model, meaning it did not give a specific value for relatedness, but instead left relatedness as an open parameter (6). In Appendix C (Model C), we ‘closed’ Model D by making additional demographic assumptions, such that a specific value for relatedness could be specified. This value of relatedness was too low for signalling to be favoured. However, this raises the question – are there any alternative ways of closing Model D, which lead to a high enough relatedness that signalling is favoured? We examine this possibility in this Appendix. Specifically, in Model C, we closed Model D by assuming that individuals signal indiscriminately to all deme-mates. In this Appendix, we close Model D by assuming, alternatively, that individuals identify their relatives and signal only to them (kin discrimination) (22).

As in our other models, we assume an infinite population of haploid, asexual individuals, split into demes of size *N* (infinite island model) (12). We focus our analysis on a focal individual, drawn at random from the population. Each generation, a random individual on each deme is initially attacked by a herbivore, and suffers a fecundity cost of *d*. Given that there are *N* individuals on each deme, each individual has a 1/*N* chance of being initially attacked by the herbivore. Individuals are each allocated a random individual from their deme to be their ‘recipient’ (individuals cannot be their own recipients).

The initially-attacked individual may then signal towards its recipient. However, signalling only occurs if the initially-attacked individual is *identical by descent* to its recipient at the signalling locus (signalling-IBD). ‘Identical by descent’ means that individuals have inherited the same allele from a shared common ancestor. (In technical population-genetic jargon, alleles are said to be ‘identical by descent’ in the infinite island model if they ‘coalesce in finite time’ (12)). If individuals are identical by descent, this means they are relatives (kin), and so by restricting signalling to identical by descent individuals, individuals are engaging in kin discrimination (22). Note that there are many other ways to recognise kin; our assumption that individuals can recognise kin based on signalling-IBD is just for illustration, chosen because it leads to fairly straightforward maths. Whether or not plants could actually recognise kin in this way (by recognising identical alleles for the social behaviour) is an open empirical question, which has recently been discussed by Montazeaud & Keller (23).

In keeping with our other models, we use the following notation. If the focal individual signals, it invests an amount *x*_*focal*_ into signal production. Signal investment incurs a fecundity cost that scales linearly with *c*. Therefore, the focal individual pays a signalling cost of *cx*_*focal*_ when it signals. The signal is transferred to the focal individual’s recipient, warning of herbivore attack. Consequently, the recipient prepares for attack, at a fecundity cost that scales linearly with *s*. We denote the average signalling investment (conditional upon the presence of a signalling-IBD recipient, resulting in signalling) amongst the *N*-1 other plants on the focal individual’s deme by *x*_*others*_. This means that, when the focal individual is signalled to by an initially-attacked individual, the focal individual will ultimately suffer a cost of attack that is given by *d(1-x*_*others*_), and a defence cost that is given by *sx*_*others*_. We assume that preparing for herbivore attack is less costly than the fecundity cost of being attacked (*d*>*s*), which ensures that individuals are favoured to defend themselves against herbivores rather than let themselves be eaten.

We emphasise that only the initially-attacked individual has a chance to warn the other individuals about the presence of a herbivore. However, as in our other models, we assume that subsequently-attacked individuals may still produce a (useless) signal, if they are unable to facultatively turn off signal production. Specifically, we assume that subsequently-attacked individuals produce a (useless) signal if their allocated ‘recipient’ from their deme is a relative (signalling-IBD), and with probability 1-*g*, where *g* denotes the extent to which signal investment is ‘facultative’ as opposed to ‘obligate’. If the focal individual produces a signal when subsequently-attacked, it pays a signalling cost of *cx*_*focal*_*(1-g)*.

After the fecundity costs of attack and signal investment have been incurred by all individuals, each individual produces a large number of offspring (juvenile haploid clones) in proportion to their fecundity. A proportion of these offspring, *m*, migrate to a different deme, randomly chosen for each juvenile individual. A proportion, *1-m*, remain on the local deme. A random sample of *N* juvenile individuals are then chosen, for each deme, to survive and form the adult population for the next generation (local population regulation). This lifecycle then iterates over many generations until an evolutionary end point is reached.

In keeping with our other models, we denote the population average investment into signalling (conditional upon the presence of a signalling-IBD recipient, resulting in signalling) by *x*_*pop*_. We denote the average signalling investment on the focal individual’s deme (conditional upon the presence of a signalling-IBD recipient, resulting in signalling) by *x*_*deme*_; Equation A1 shows how *x*_*deme*_ can be written in terms of *x*_*focal*_ and *x*_*others*_. As usual, we assume that the baseline fecundity is 1, that 0 ≤ *x*_*focal*_, *x*_*others*_, *x*_*deme*_, *x*_*pop*_ ≤ 1, and that *c* + *d* + *s* ≤ 1.

The probability that a given individual is signalling-IBD with its allocated recipient, in the evolutionary long-term, is given by *r*\* in Equation A16. In other words, ‘signalling-IBD’ in this model is equal to ‘relatedness’ in our previous closed Models (A & C). The reason for this equivalence is that, in our models, relatedness can be defined as the probability of being identical by descent with one’s social partner, measured at the locus responsible for encoding the social behaviour (signalling) (24). In our previous closed Models (A & C), social partners are random deme-mates. Therefore, in these Models (A & C), relatedness is given by signalling-IBD. In our present model, relatedness is *not* given by signalling-IBD (we derive relatedness in our present model below; see Equation E7). We therefore do not use the notation *r*\* for signalling-IBD in this model; instead we use *F*, which can be written explicitly as

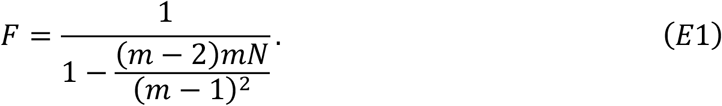

We reiterate that *F* in Equation E1 has the same value as *r*\* in Equation A16, and is derived in the same way. Readers interested in the derivation of *F* should therefore consult Appendix A to see how *r*\* in Equation A16 was derived.

Using Equation E1, we can write down the fecundity functions that will arise in the evolutionary long-term. The fecundity of the focal individual in the evolutionary long-term is given by *f*_*focal*_; the average fecundity on the focal individual’s deme in the evolutionary long-term is given by *f*_*deme*_; the average fecundity in the population in the evolutionary long-term is given by *f*_*pop*_. These fecundity functions can be written explicitly as

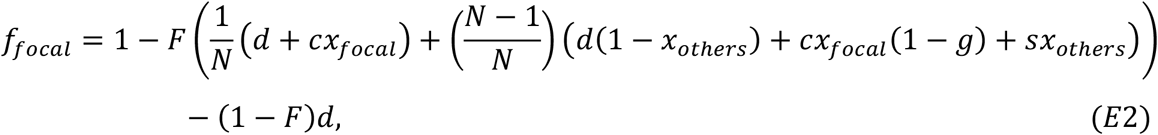

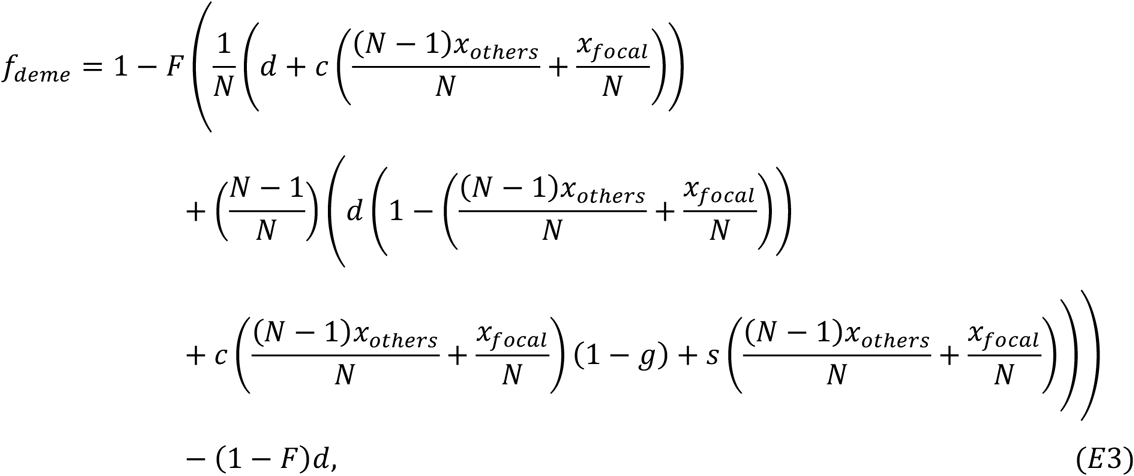

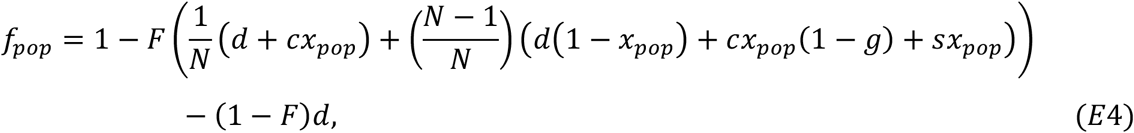

where the explicit expression for *F* is provided in Equation E1. To interpret these fecundity functions, note that they are obtained from the fecundity functions in Models A–D (Equations A2, A4 & A5) by first weighting all fitness effects by *F*. This step is taken because signalling now only occurs in a proportion of generations given by *F*, whereas in Models A–D signalling occurred every generation. Next, an extra term −(1 − *F*)*d* is added to each fecundity function, which accounts for the proportion of generations 1 − *F* in which signalling does not occur, resulting in all individuals on the deme getting attacked at a fecundity cost *d*.

We can then obtain an expression for the fitness of the focal individual (*w*) by substituting Equations E2–4 into Equation C1. To obtain the equilibrium level of signal production, *x\**, we first evaluate the following partial derivatives of the fitness function (*w*):

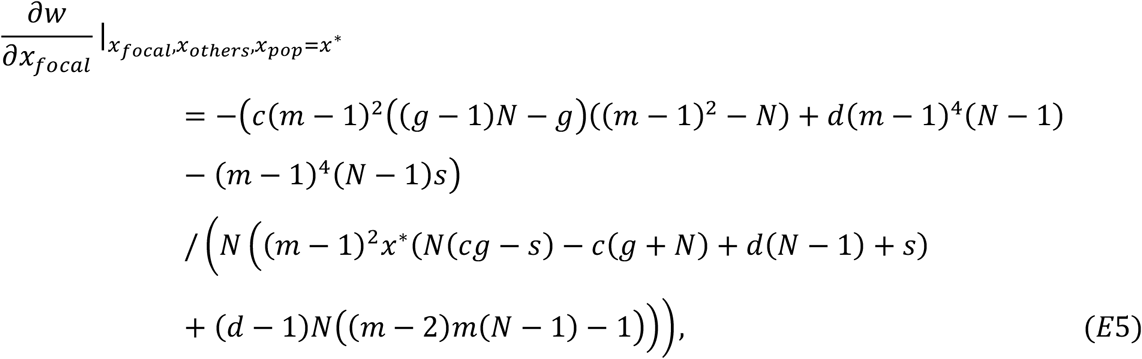

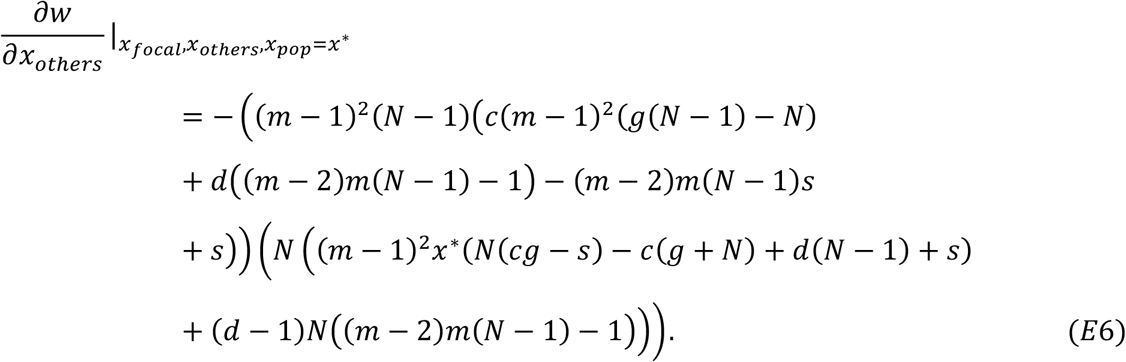

We then write down relatedness as

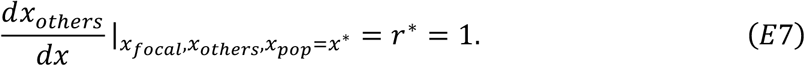

The reason why relatedness is 1 (*i*.*e*., maximal) in this model is that individuals only socially interact with (*i*.*e*., signal to) individuals that are identical by descent at the locus for the social behaviour (signalling). Therefore, any change in signal investment by an individual will result in an identical change in the signal investment of its social partners, resulting in 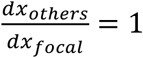 = 1 and therefore Equation E7.

Substituting Equations E5–E7, as well as the trivial partial derivative 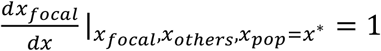, into Equation A7, we obtain the selection differential for signalling:

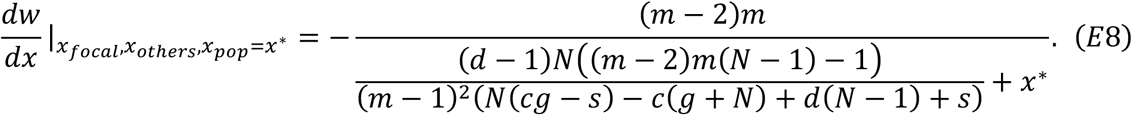

This selection differential is positive, resulting in signalling (*x*^*^ = 1), when the following condition is satisfied:

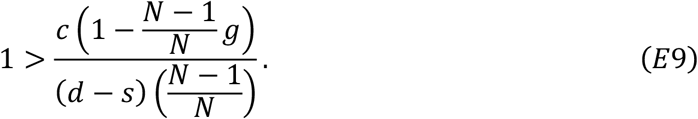

Conversely, the selection differential is negative, resulting in no signalling (*x\**=0), when 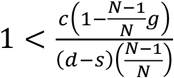 is satisfied.

Note that Equation E9 can be obtained from Equation D4 by substituting in *r*\*=1. This provides a robustness check on our open and closed models. Specifically, Equation D4 gives the condition for signalling to be favoured, in an ‘open’ model with migration (*m*=1); it states that signalling is favoured if relatedness (*r*\*) is high enough relative to other model parameters. Equation E9 gives the condition for signalling to be favoured, in a ‘closed’ model where *r*\*=1 due to kin discrimination. The latter condition (Equation E9) is obtained by substituting the specific value for relatedness in this closed model into Equation D4 (*r*\*=1).

We explained in Appendix D that Equation D4 is a form of Hamilton’s rule. Therefore, given that Equation E9 is obtained from Equation D4 by specifying a specific relatedness value (*r*\*=1), we can say that Equation E9 is a form of Hamilton’s rule, albeit one where the relatedness parameter is fixed. We can see this by setting 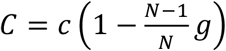, 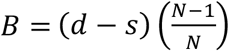 and *r*\*=1 to obtain 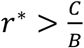. The capital ‘C’ gives the overall cost of the social act (signalling). It is the unit cost of signal production (*c*), weighted by incidences where this cost is actually paid 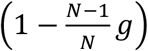. The capital ‘B’ gives the overall benefit of the social act (signalling). It is the unit benefit of signal production, as manifested as a reduced fecundity cost of being attacked (*d*) minus the cost of herbivore defence (*s*), weighted by incidences where this benefit is actually received 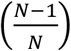.

Overall, signalling can evolve in this closed model with kin discrimination. This contrasts with Appendix C (Model C), which is an equivalent model except that individuals signal indiscriminately. Therefore, kin discrimination facilitates the evolution of signalling.

## Appendix F

### Model F: Inclusive fitness analyses

Our analyses so far have used ‘neighbour-modulated fitness’ rather than ‘inclusive fitness’. The alternative fitness measures differ in the way they count up fitness effects (1). ‘Neighbour-modulated fitness’ counts up all fitness effects *experienced by* a focal individual due to its social environment. ‘Inclusive fitness’ counts up all fitness effects *caused by* a focal individual due to its social behaviour. Under a standard set of mathematical assumptions (*e*.*g*., weak selection), the two approaches give the same answer; *i*.*e*., they predict the same optimised trait values (*x*\*) (2, 12, 14–20). However, some people find inclusive fitness analyses more intuitive and easier to follow (5), so in that spirit, we present inclusive fitness analyses of Models C & E. We have chosen these models because they make the same lifecycle assumptions except for one difference: in Model C, individuals signal indiscriminately, whereas in Model E, individuals signal discriminatorily towards kin. Signalling evolves in the ‘kin discrimination’ scenario, but not in the ‘indiscriminate signalling’ scenario. Our inclusive fitness analyses will illustrate why signalling only evolves in one of these scenarios. For concision, we do not present inclusive fitness analyses of all our Models, but these could be obtained by following an analogous approach to the one we will outline in this Appendix.

In Models C & E, a signalling investment of *x* results in a cost to the signaller of *C* and a benefit to the signal receiver of B (where *B & C* are functions of *x*). We can provide explicit expressions for *B* and *C*, but to keep things general for now, we leave them unspecified. Signalling therefore adds *B–C* offspring to the population, and a proportion of these 1–*m* will remain on the natal deme. Given that deme size is constant, this means that an act of signalling results in (1–*m*)(*B–C*) offspring being displaced from the natal patch.

Relatedness between an actor (signaller) and recipient (signal receiver) is denoted by *r*. Relatedness between an actor and a random member of its deme is denoted by *r*_*whole*_. Relatedness between an actor and the individuals who are competitively displaced as a result of its signalling is given by (1–*m*)*r*_*whole*_ (the reason why the 1-*m* factor is applied is that a proportion of displaced offspring *m* will have immigrated in from other demes and so will be related by zero). We calculate the marginal inclusive fitness of signalling by summing up all relatedness-weighted fitness effects arising from an act of signalling:

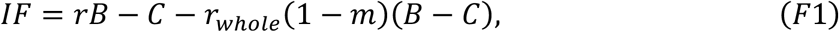

where *B* and *C* are functions of signalling investment, *x*.

We have already calculated explicit expressions for *r*_*deme*_ and *r*. When individuals signal indiscriminately, *r* is given by Equation A16 and *r*_*whole*_ is given by Equation A14. Substituting these explicit expressions into Equation F1, we find that *IF*>0 is never satisfied. Consequently, indiscriminate signalling is not favoured. This recovers the result of our ‘neighbour-modulated fitness’ analysis presented in Appendix C. Note that we obtained this result without specifying values for *B* and *C*. This clarifies that, for this lifecycle, indiscriminate helping (which indiscriminate signalling is an example of) is never favoured, irrespective of the explicit expressions for *B* and *C*. This is a well-established result in the social evolution literature (7, 11, 12).

When individuals signal discriminatorily towards kin, *r* is given by Equation E7 (*i*.*e*., *r*=1). *r*_*whole*_ is given by Equation A14, as was also the case in the indiscriminate signalling scenario. Substituting these explicit expressions into Equation F1, we find that *IF*>0 is satisfied when

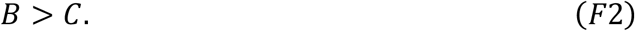

In other words, given that relatedness between social partners is maximal (*r*=1), signalling is favoured if the benefit to the signal-receiver is greater than the cost to the signaller. We can write down explicit functions for *B* and *C* for our model:

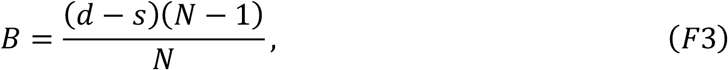

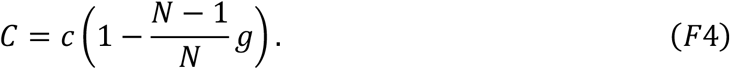

*C* is obtained by multiplying the unit cost of signal production (*c*) by the proportion of generations where this cost is actually paid 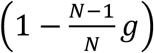. *B* is obtained by multiplying the net fecundity gain of being warned about a herbivore (*d*–*s*) by the proportion of generations where this benefit is actually given 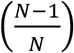. Substituting Equations F3 & F4 into Equation F2, we find that *IF*>0 holds when Equation E9 is satisfied. Therefore, we exactly recover the result of Appendix E, this time using an inclusive fitness rather than neighbour-modulated fitness analysis. Overall, we can see that kin discrimination can facilitate the evolution of signalling.

## Appendix G

### Model G: Cue-suppression; closed; population regulation before migration

We now slightly alter assumptions of Model A, to study a different sort of social behaviour. As before, we assume an infinite population of haploid and asexual individuals, split into demes of size *N*, and focus on a randomly drawn focal individual. As before, each generation, a random individual on each deme is initially attacked by a herbivore and suffers a fecundity cost *d*.

However, in the present model, after the initially-attacked individual is attacked by the herbivore, it releases a *cue* rather than a *signal*. The cue indicates to the other *N*-1 individuals on the deme that a herbivore is in the vicinity. However, we assume that the initially-attacked individual may suppress the cue, to stop the other *N*-1 individuals from learning that a herbivore is present. If the focal individual is the initially-attacked individual on its deme, it invests an amount *x*_*focal*_ into cue suppression. Investment into cue-suppression incurs a fecundity cost that scales linearly with *c*. Therefore, the focal individual pays a cue-suppression cost of *cx*_*focal*_ when it is initially-attacked.

We denote the average cue-suppression investment amongst the *N*-1 other plants on the focal individual’s deme by *x*_*others*_. This means that, when the focal individual is not the initially-attacked individual, it suffers a cost of attack of *dx*_*others*_, and a defence cost that is given by *s(1-x*_*others*_). We denote the population average investment into cue-suppression by *x*_*pop*_, and the average cue-suppression investment on the focal individual’s deme by *x*_*deme*_.

As before, we assume that only the initially-attacked individual can convey useful information about herbivore presence. Specifically, only cues transmitted by the initially-attacked individual can warn the other individuals about herbivore-presence; cues transmitted by subsequently-attacked individuals are useless, because the herbivore has already arrived. We allow for the facultative ‘turning off’ of cue-suppression when the individual is subsequently-attacked. Therefore, the focal individual pays a cue-suppression cost of *cx*_*focal*_*(1-g)* when subsequently-attacked (non-focal individuals pay a signalling cost of *cx*_*others*_*(1-g)* when subsequently-attacked), where *g* denotes how facultative cue-suppression is.

After social interactions and attacks have taken place, the rest of the lifecycle plays out as it does in the Model A: production of a large number of juvenile offspring in proportion to fecundity; random sampling of *N* juveniles on each deme to establish the next adult population; migration of (surviving) adults to a random patch with probability *m*. The lifecycle is iterated until an evolutionary end-point is reached.

We emphasise that the only mathematical difference between this model and Model A is that, in Model A, secondarily-attacked individuals suffer attack and defence costs of *d*(1-*x*_*i*_) & *sx*_*i*_ respectively, but in the present model, secondarily-attacked individuals suffer attack and defence costs of *dx*_*i*_ *& s*(*1-x*_*i*_) respectively (where *i*=*others* for the focal individual; *i*=*deme* for non-focal individuals on the focal individual’s deme; *i*=*pop* for individuals on other demes). The other mathematical details are the same. This means the fecundity functions (Equations A2–A5) are changed to:

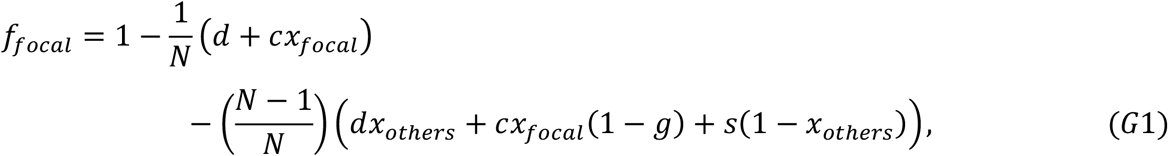

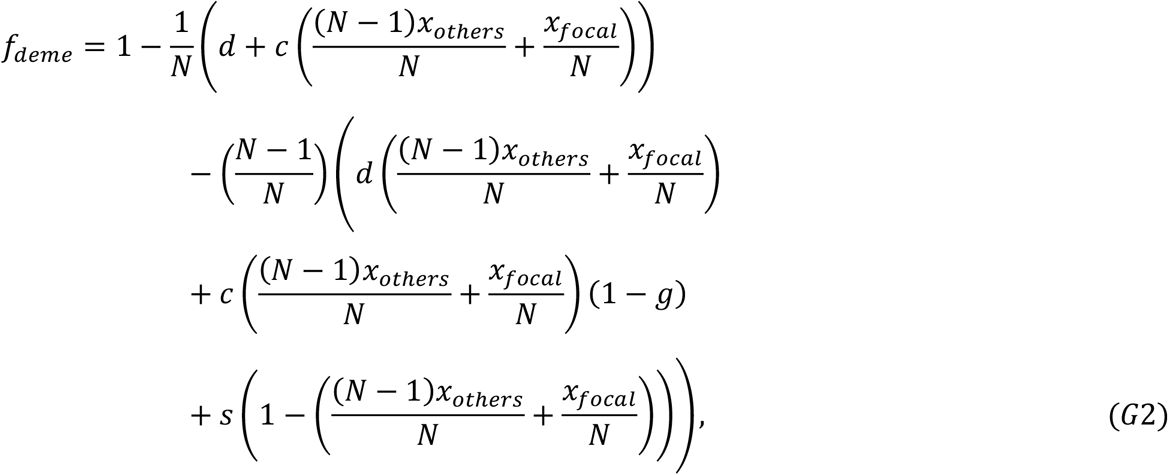

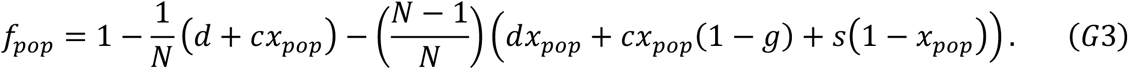

The fitness of the focal individual is obtained by substituting the fecundity functions (Equations G1 – G3) into Equation A6. With fitness obtained, we proceed, as usual, by evaluating the derivatives on the right-hand side of Equation A7, which we now obtain as

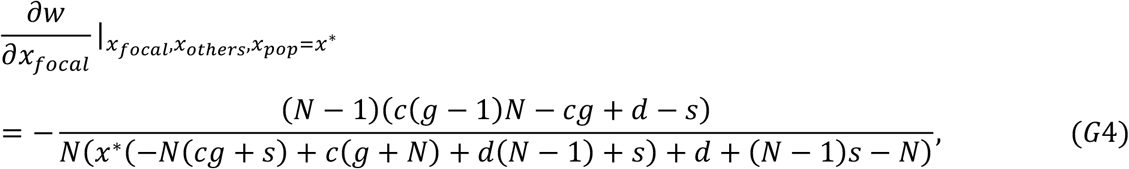

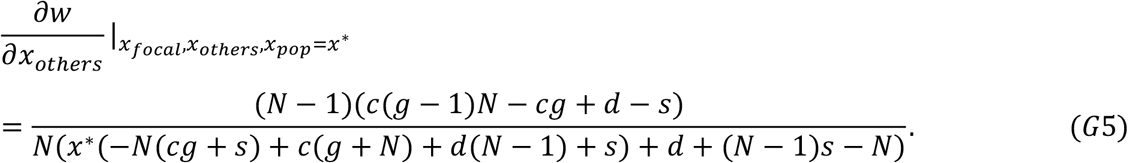

We obtain the same expressions for 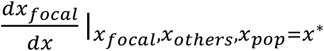 and 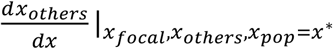 as we did for Model A (provided in Equations A8 and A17 respectively). By substituting Equations A8, A17, G4 & G5 into Equation A7, we obtain

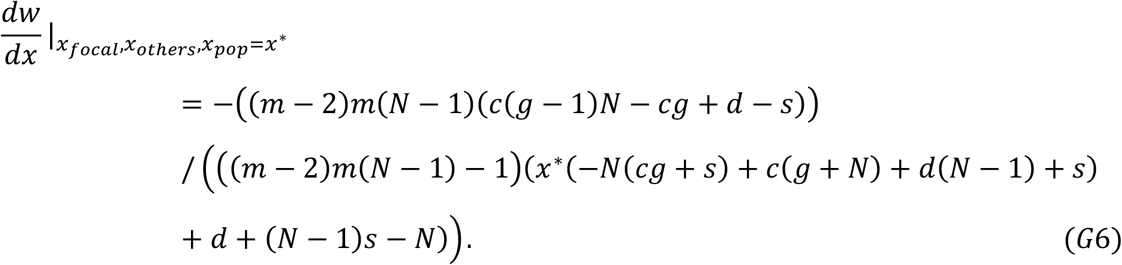

We can see by setting this to be greater than zero that cue-suppression evolves when

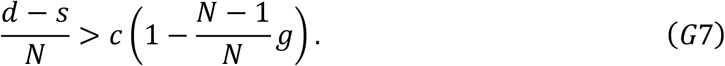

Equation G7 can be understood as cue-suppression (harming) being favoured when the cost of the social act to the actor, 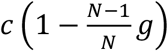, is less than the cost of the social act to the competitor, 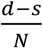. The actor cost term 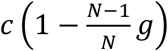 is the unit cost of cue-suppression (*c*), weighted by the proportion of incidences where this cost is actually paid 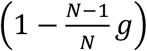. The competitor cost term 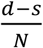 is the fecundity cost of being attacked (*d*), minus the saved cost of herbivore defence (*s*), weighted by the proportion of incidences where these costs can actually be applied by the actor (1/*N*).

Therefore, for this lifecycle (population regulation before migration), suppressing a cue of herbivore-presence can evolve, but signalling herbivore-presence cannot evolve (Model A).

## Appendix H

### Model H: Cue-suppression; open; population regulation before migration

In this Appendix, we ‘open’ Model G, by substituting Equation A12 rather than Equation A17 into Equation A7 to obtain

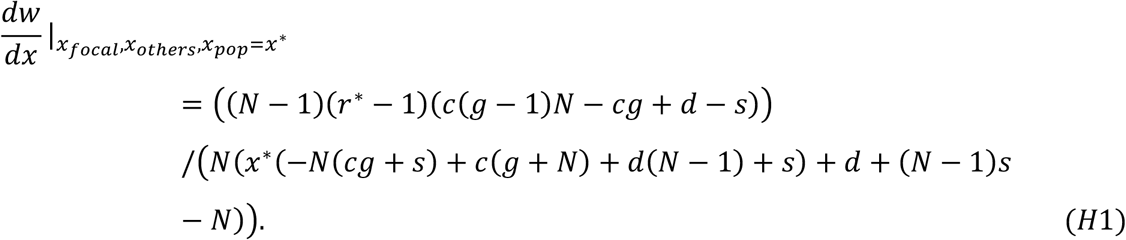

We can see by setting this to be greater than zero that cue-suppression evolves when Equation G7 is satisfied 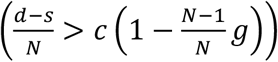. The conditions are the same for the open and closed versions of the model, which shows that allowing relatedness to vary independently of kin competition does not affect evolutionary outcomes (relatedness does not influence the outcome of social evolution in this lifecycle).

## Appendix I

### Model I: Cue-suppression; closed; population regulation after migration

We now amend Model G so that population occurs after migration, rather than before. This means that fitness is obtained by substituting the fecundity functions Equations G1 – G3 into Equation C1 rather than A6, leading to the following derivatives:

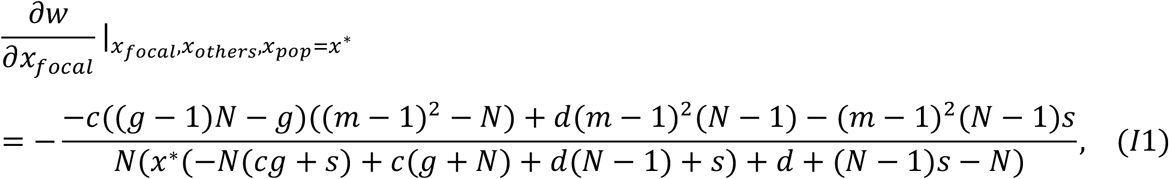

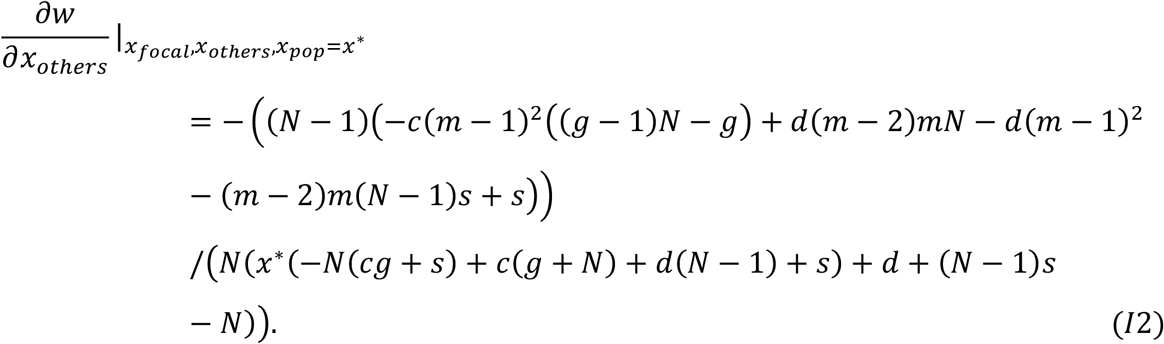

We obtain the same expressions for 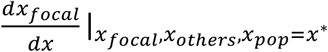 and 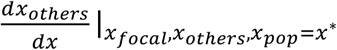 as we did for Model A (provided in Equations A8 and A17 respectively).

By substituting Equations A8, A17, I1 & I2 into Equation C1, we obtain

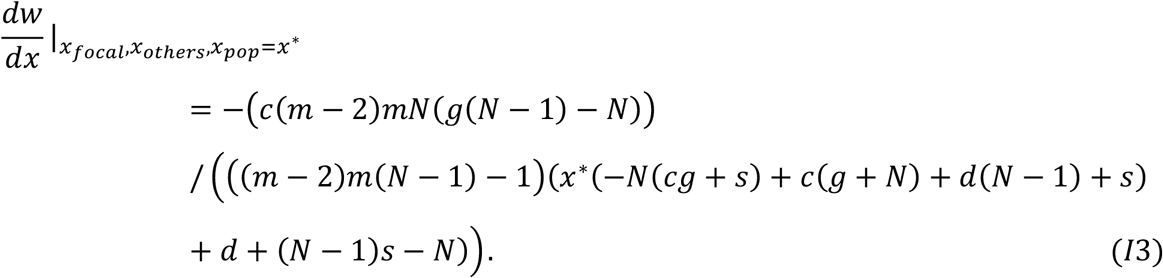

As long as there is some migration (*m*>0) and cue-suppression cost (*c*>0), the right-hand side of Equation I3 is strictly negative. This means that the long-term (ESS) level of cue-suppression is *x*^*^ = 0. Therefore, at equilibrium, individuals evolve to *not* suppress the cue of herbivore-presence. Any tendency to suppress the cue of herbivore-presence will ultimately be removed by natural selection. Therefore, cue-suppression could evolve previously (Models G & H), but the change in lifecycle assumption, so that population regulation occurs after migration rather than before, prevents cue-suppression from evolving.

## Appendix J

### Model J: Cue-suppression; open; population regulation after migration

We now ‘open’ Model I, by substituting Equation A12 rather than Equation A17 into Equation C1, to obtain

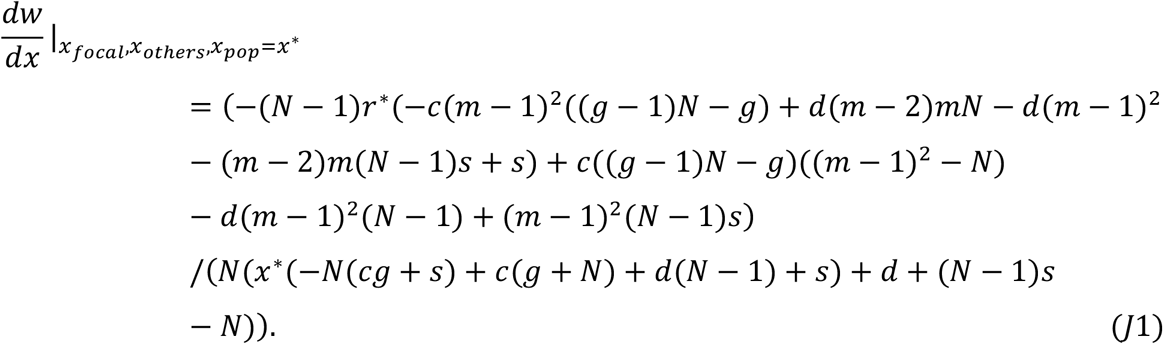

This expression is complex, so we focus on two special cases. First, we consider the case where the migration rate is one. Substituting *m*=1 into Equation J1, we obtain

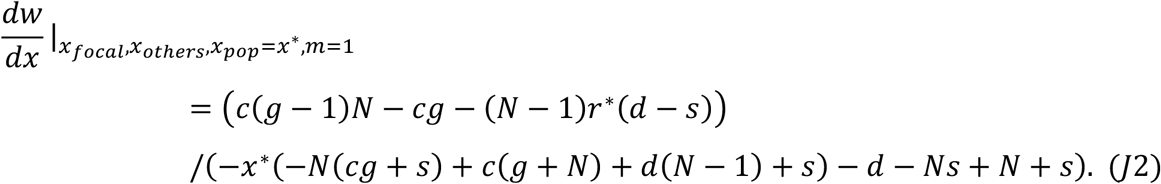

As long as there is some relatedness (r*>0) or cost of cue-suppression (*c*>0), the right-hand side of Equation J2 is strictly negative. This means that, when there is maximal migration, the long-term (ESS) level of signal production is *x*^*^ = 0 (no cue suppression).

Second, we consider the case where the migration rate is zero. Substituting *m*=0 into Equation J1, we obtain Equation H1, which is positive (meaning cue-suppression is favoured, leading to *x*\*=1) whenever Equation G7 is satisfied 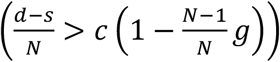. Conversely, cue-suppression is disfavoured, leading to *x\**=0, whenever 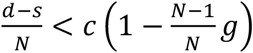. Therefore, cue-suppression is sometimes favoured if there is sufficiently low migration (*m*).

Therefore, whilst cue-suppression could evolve when population regulation preceded migration (Models G & H), but not in a closed model where migration preceded population regulation (Model I), opening this latter model (I) allows the results of the previous models (G & H) to be recovered for the zero-migration case.

## Appendix K

### Supplementary Discussion

#### Summary of results

To recap we found that, when relatedness is detached from model parameters (open model) and sufficiently high that Hamilton’s Rule is satisfied (Equation D4), and migration precedes population regulation and is sufficiently high, signalling can evolve (Appendix D). However we found that, when relatedness emerges from model parameters (closed model), relatedness is only high enough for signalling to be favoured if individuals are capable of identifying and signalling towards relatives (kin discrimination), rather than signalling indiscriminately.

We found that suppressing a cue of herbivore-presence is favoured when cue-suppression causes more harm to others than yourself (Equation G7), and either: population regulation precedes migration (Models G & H); or, migration precedes population regulation, relatedness is detached from model parameters and sufficiently low, migration is sufficiently low (Model J).

To explain these results, we first note that: signalling the presence of a herbivore is an example of a *helping* behaviour because it increases the fitness (offspring after one lifecycle iteration) of neighbours; suppressing a cue of herbivore-presence is an example of a *harming* behaviour because it decreases the fitness of neighbours (7, 12, 25). Most of our results are recovered as special cases of previously published results on the evolution of helping and harming behaviours generally (though this doesn’t render our theoretical analysis redundant or trivial – our models had novel ecological aspects like herbivore attack, which could in principle have caused our results to diverge from previous models).

#### Models C & I

As first shown formally by Taylor (10), in closed island lifecycles where population regulation goes after migration, indiscriminate social behaviours (i.e., helping and harming behaviours) cannot evolve. (Technically, indiscriminate behaviours with social characteristics can evolve, but only if they provide a ‘direct’ fitness benefit to the actor – in other words, the ‘indirect’ or ‘social’ characteristic of the behaviour does not have any bearing on whether it is favoured; 7.) This is why neither signalling herbivore-presence nor suppressing a cue of herbivore-presence could evolve in our Models C & I. The reason for this is that, in closed models of this type, indiscriminate social behaviours either serve to help competitors, or harm non-competitors, which precludes the evolution of social behaviours.

To tease this out further, consider an indiscriminate helping behaviour. For helping to evolve (by kin selection), help needs to be directed towards individuals that are: (*i*) sufficiently related to the actor, and (*ii*) in sufficiently weak competition with the actor (5, 7–11). But the closed lifecycle means that, whenever help is directed towards relatives (this occurs when the migration rate is low), the individuals receiving help are also in strong competition with the helper (because the low migration rate leads to intense local competition). There are no combinations of demographic parameters (*N,m*) that simultaneously lead to high relatedness and low competitiveness amongst social partners, which precludes the evolution of indiscriminate helping.

Now consider an indiscriminate harming behaviour. For indiscriminate harming to evolve, harm needs to be directed towards individuals that: (*i*) have sufficiently low relatedness with the actor; (*ii*) are in sufficiently strong competition with the actor (7, 9, 26, 27). But the closed lifecycle means that, whenever harm is directed towards non-kin (this occurs when the migration rate is high), the individuals that are harmed are not in strong competition with the harmer (because the high migration rate causes them to compete appreciably on other demes). There are no combinations of demographic parameters (*N,m*) that simultaneously lead to high competitiveness and low relatedness amongst social partners, which precludes the evolution of indiscriminate harming.

We note that these results hold irrespective of the fitness of the actor, and are supported by a large theoretical literature stemming from Taylor’s (10) pivotal paper. This rules out, for instance, ‘death screams’ (28), whereby a plant that is about to be fatally eaten releases one final warning signal before dying. ‘Death screams’ are disfavoured, even as a ‘best of a bad job’, because all individuals on the deme would hear this scream, leading to increased competition, and so no individual on the patch would gain an overall competitive benefit.

#### Models D & J

Taylor’s (10) scenario can be ‘opened’ so that relatedness can vary independently of demographic parameters (*N,m*). Consequently, relatedness can be made to increase, without also increasing the relatedness of competitors, allowing indiscriminate helping behaviours to evolve. Specifically, this requires that, as well as a high value for the relatedness parameter, the migration rate is kept high, leading to low competition between the actor and its social partners. Frank (4, p114-115) showed that increasing a relatedness parameter, as well as a migration parameter (or in his terminology, a ‘spatial scale of density dependent competition’ parameter), can allow indiscriminate helping to evolve. This explains why signalling herbivore-presence (an example of a helping behaviour) could evolve in our Model D.

Analogously, ‘opening’ the Taylor (10) scenario can allow indiscriminate harming behaviours to evolve, by simultaneously decreasing a relatedness parameter, as well as a migration parameter (increasing competitiveness with social partners). This explains why cue-suppression (an example of a harming behaviour) could evolve in our Model J.

#### Model E

Taylor’s (10) scenario can be amended so that individuals socially interact preferentially with relatives (kin discrimination), as opposed to interacting randomly with deme-mates (indiscriminate helping). This causes relatedness towards social partners to increase, without also increasing relatedness towards competitors (deme-mates), allowing helping behaviours to evolve. This result has been obtained numerous times, for a variety of kin recognition mechanisms (29–31). This explains why signalling could evolve in our kin discrimination model (Model E).

#### Models A & G

Taylor’s closed (10) scenario can be amended slightly, so that population regulation precedes migration. Lehmann and Rousset (7) showed that helping behaviours cannot evolve in this setting, but harming behaviours can. This is why signalling herbivore-presence (a helping behaviour) could not evolve in our Model A, but cue-suppression (a harming behaviour) could evolve in our Model G.

The reason why helping cannot evolve in this lifecycle is that, when population regulation precedes migration, competition occurs completely within demes (rather than between individuals from different demes), leading to maximally local competition. The evolution of helping requires that competitiveness with social partners is low, but this is impossible in this lifecycle (i.e., it cannot be achieved by any combination of demographic parameters, *N & m*), precluding the evolution of helping.

The reason why harming can evolve in this lifecycle is that, for harming to evolve, there needs to be sufficiently high competition and low relatedness with social partners. In this lifecycle, competition with social partners is maximal (maximally local competition). It turns out that, given the extreme strength of local competition, harming can evolve for any value of relatedness (*i*.*e*., it doesn’t need to be especially low, because local competition is so intense) (7).

#### Models B & H

Lehmann and Rousset’s (7) closed scenario, where population regulation precedes migration, can be ‘opened’ so that relatedness can vary independently of demographic parameters (*N,m*). However, opening this scenario (*e*.*g*., Models B & H) does not alter any of the equivalent closed model results (*e*.*g*., Models A & G). Specifically, helping can still never evolve, and harming can (for any relatedness). The reason for this is that, in the open model setting, relatedness to social partners can be varied, but this is inconsequential, because it is the intense (maximal) local competition that is driving results for this lifecycle. Specifically, it precludes helping behaviours, and facilitates harming behaviours, irrespective of relatedness to social partners.

#### Take-home message

Closed models are more realistic than open ones, because they give a full account of how relatedness emerges from demography. Open models can be useful, particularly as a way of interpreting their counterpart closed models, by detaching relatedness from model parameters, so that the causal effect of kin selection on model outcomes can be isolated and examined (4, 6).

In the closed models considered, suppressing a cue of herbivore-presence could evolve (if population regulation precedes migration), but signalling herbivore-presence could only evolve if individuals can identify and signal preferentially towards (kin discrimination). The take-home message is therefore that suppressing a cue of herbivore-presence is likely to evolve more permissively than signalling herbivore-presence, unless plants are capable of widespread kin discrimination.

## Appendix L

### Model L: signalling; open & closed; population regulation before migration; individuals may signal dishonestly and not respond to signals

In this appendix, we generalise Model A, so that individuals can choose to signal dishonestly (*i*.*e*., warn others of herbivore-presence when the herbivore isn’t there), and choose not to respond to the signal (*i*.*e*., not prepare for herbivore-attack when warned by another individual). We have written the verbal model description for this appendix entry in full, without referring to previous appendices, so that it can be read in isolation. In this appendix, we examine both ‘closed’ (emergent relatedness) and ‘open’ (parameterised relatedness) versions of the model, starting with the closed version.

Each generation, with probability *p*, the population is attacked by herbivores (with probability 1-*p*, the population is not attacked). In generations where the population is attacked, a random individual on each deme is initially attacked by a herbivore, and suffers a fecundity cost of *d*. Given that there are *N* individuals on each deme, each individual has a 1/*N* chance of being initially attacked by the herbivore. The initially-attacked individual may then invest into the production of a signal. If the focal individual is the initially-attacked individual on its deme, it invests an amount *x*_*focal*_ into signal production. Signal investment incurs a fecundity cost that scales linearly with *c*. Therefore, the focal individual pays a signalling cost of *cx*_*focal*_ when it is initially-attacked.

The signal produced by the initially-attacked individual is transferred to (and recognised by) the other *N*-1 individuals on the deme. We make no assumptions about how the signal is transferred, but it may be being transferred through a common mycorrhizal network, or through the air, *etc*. The signal warns the other plants that a herbivore is in the vicinity. Consequently, the other plants can prepare for being attacked, and defend themselves, meaning they can reduce their fecundity cost of attack. Plants may choose the extent to which they actually respond to the signal, preparing for herbivore-attack, rather than ignoring the signal. Preparation for herbivore attack (defence) incurs a fecundity cost that scales linearly with *s*. We denote the average signalling investment amongst the *N*-1 other plants on the focal individual’s deme by *x*_*others*_. We denote the extent of signal-response by the focal individual by *y*_*focal*_. This means that, when the focal individual is not the initially-attacked individual, it will ultimately suffer a cost of attack that is given by *d(1-x*_*others*_*y*_*focal*_), and a defence cost that is given by *sx*_*others*_*y*_*focal*_. We assume that preparing for herbivore attack is less costly than the fecundity cost of being attacked (*d*>*s*), which ensures that individuals are favoured to defend themselves against herbivores.

In generations where the population is not attacked by herbivores (such generations occur with probability 1-*p*), a random individual on each deme is given the opportunity to feign being an initially-attacked individual, producing the signal of herbivore-presence even though no herbivore is present. In such cases, any signal produced is ‘dishonest’. If the focal individual is given the opportunity to be the dishonest signaller, it invests *cx*_*focal*_*z*_*focal*_ into signal production, where *z*_*focal*_ denotes dishonesty, with *z*_*focal*_=0 corresponding to no dishonest signal production, and *z*_*focal*_=1 corresponding to the same investment in signal production irrespective of whether the herbivore is present or not. If the *focal* individual is not given the opportunity to be the dishonest signaller, it will suffer a cost of preparation for herbivore attack that is given by *sx*_*others*_*y*_*focal*_*z*_*others*_, and a cost of obligate (redundant) dishonest signal production of *cx*_*focal*_*(1-g)z*_*focal*_.

We denote the population average investment into signalling by *x*_*pop*_. We denote the average signalling investment on the focal individual’s deme (incorporating its own signalling investment) by *x*_*deme*_; this can be written in terms of *x*_*focal*_ and *x*_*others*_ as 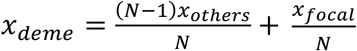 (Equation A1). Similarly, we denote the population average investment into signal response and dishonesty, respectively, by *y*_*pop &*_ *z*_*pop*_, and the *pop*-deme-average investment into signal response and dishonesty, respectively, by *y*_*deme &*_ *z*_*deme*_, where

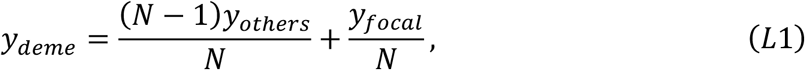

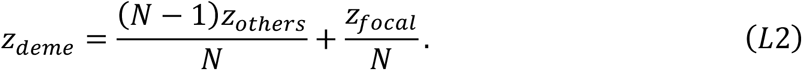

We assume that the baseline fecundity is 1, that 0 ≤ *x*_*focal*_, *x*_*others*_, *x*_*deme*_, *x*_*pop*_, *y*_*focal*_, *y*_*others*_, *y*_*deme*_, y_*pop*_, *z*_*focal*,_ *z*_*others*_, *z*_*deme*_, *z*_*pop*_ ≤ 1, and that *c* + *d* + *s* ≤ 1; these assumptions ensure that fecundity never falls below zero. The number of juvenile offspring produced by the focal individual is then proportional to

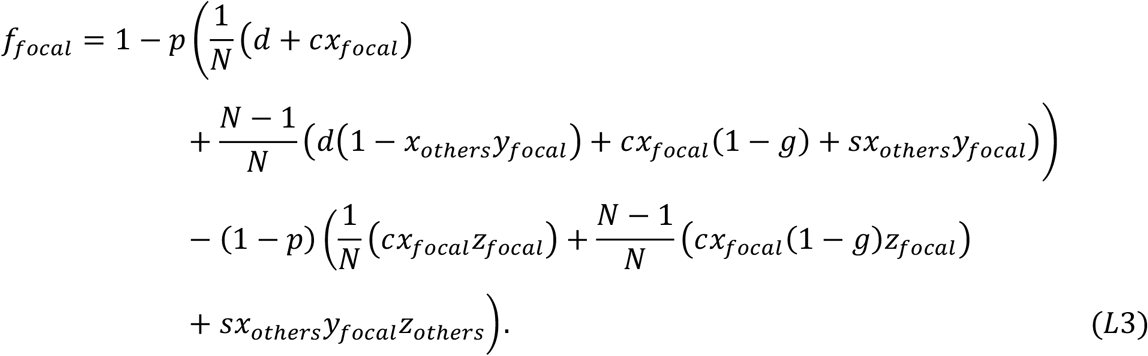

We can make sense of the right-hand side of Equation L3 as follows. Baseline fecundity is one (left-hand term). With probability *p*, a herbivore attacks. Given herbivore-attack, with probability 1/*N*, the focal individual is attacked first, leading to additively applied costs of attack (*d*) and signal production (*cx*_*focal*_). Given herbivore-attack, with probability 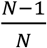, the focal individual is not attacked first, and can receive a signal of herbivore presence from another individual on the deme, leading to additively applied costs of attack (*d*(*1-x*_*others*_)), defence (*sx*_*others*_*y*_*focal*_), and the costs of signal production that are incurred even when the focal individual is not attacked first, owing to obligatory (unconditional) expression (*cx*_*focal*_(*1-g*)). With probability 1-*p*, a herbivore does not attack. Given herbivore-absence, with probability 1/*N*, the focal individual is given the opportunity to feign herbivore-presence, leading to an additively applied cost of dishonest signal production (*cx*_*focal*_*z*_*focal*_). Given herbivore-absence, with probability 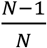, the focal individual is not given the opportunity to feign herbivore-presence, and can receive a dishonest signal of herbivore presence from another individual on the deme, leading to additively applied costs of defence (*sx*_*others*_*y*_*focal*_*z*_*others*_), and the costs of dishonest signal production that are incurred even when the focal individual is not given the opportunity to feign herbivore-presence, owing to obligatory (unconditional) expression (*cx*_*focal*_(*1-g*)*z*_*focal*_).

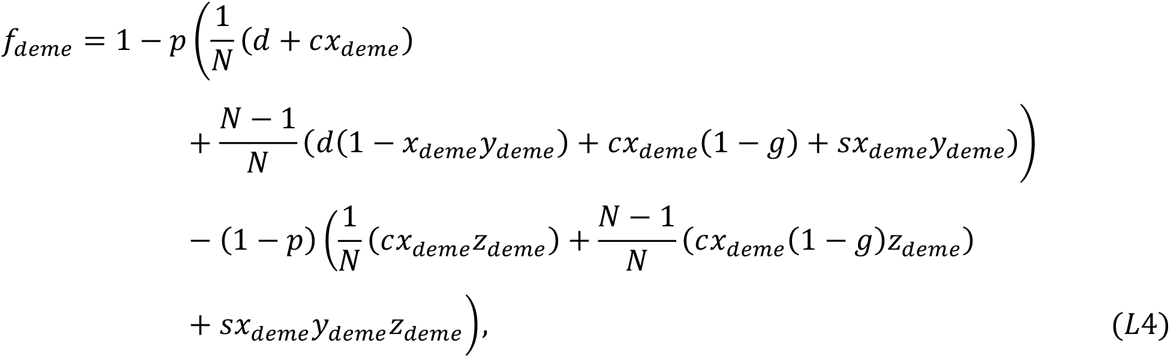

which we can rewrite using Equations A1, L1 & L2 (getting rid of the superfluous *x*_*deme*_, *y*_*deme*_& *z*_*deme*_ parameters) as

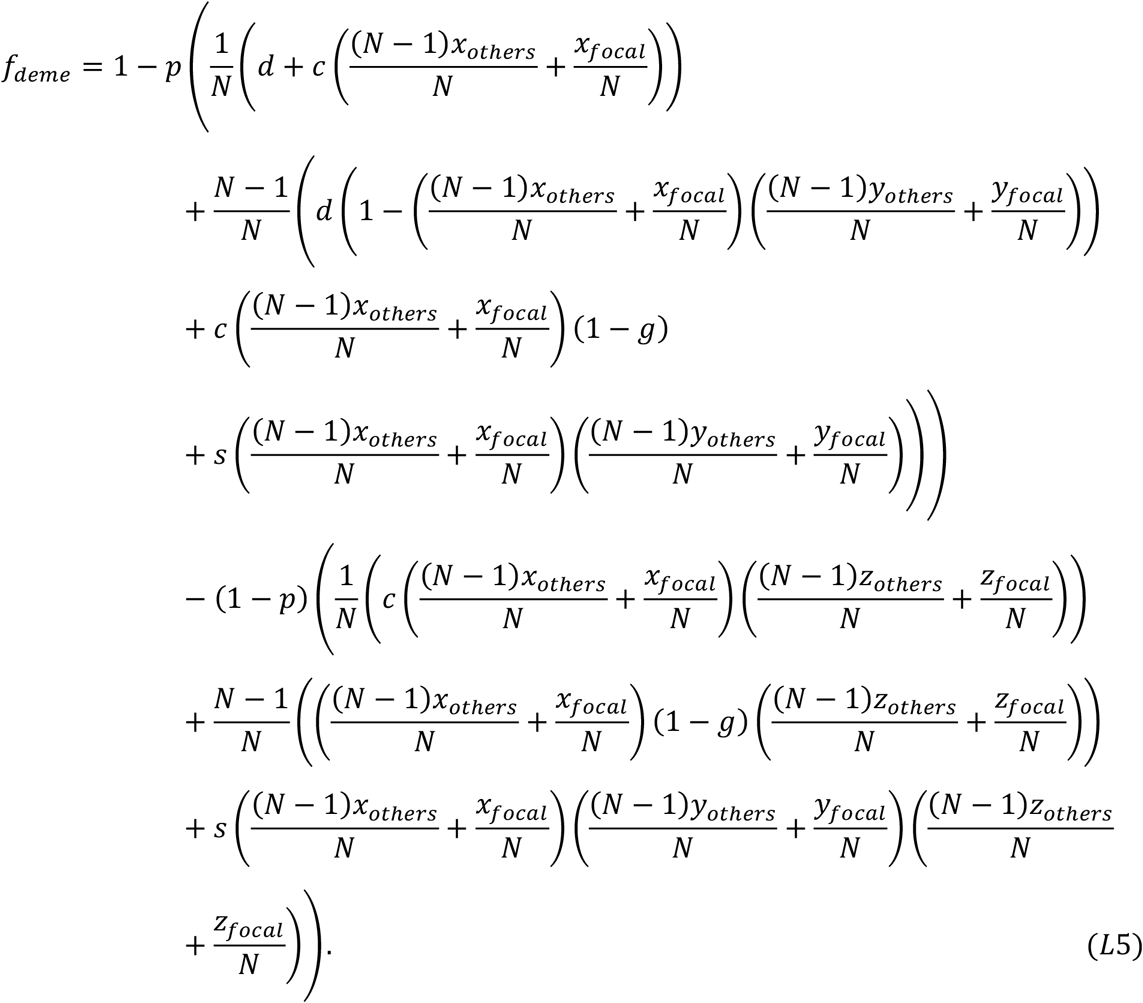

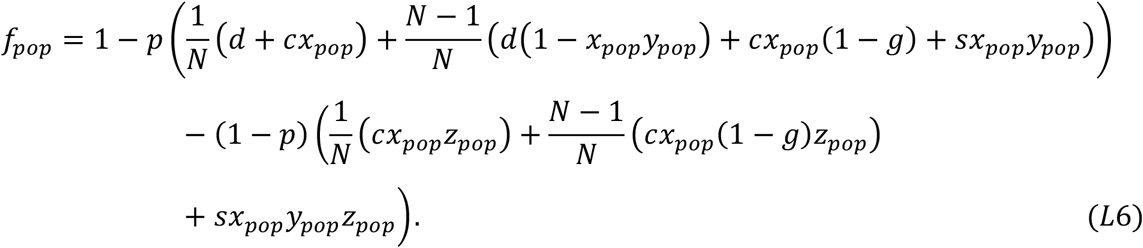

The fitness of the focal individual is then given by Equation A6 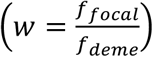, where explicit expressions for the fecundity (*f*_*focal*_, *f*_*deme*_) functions are provided in Equations L3 & L5 (7, 12, 13). Note that the *f*_*pop*_ function derived above (Equation L6) does not feature in this fitness function, but it will feature in the fitness function of subsequent models. Equation A6 reflects the fact that, under our assumption that (local) population regulation occurs before migration, individuals only compete with competitors on their native deme, rather than with individuals on other demes (see Appendix A for more discussion).

Our aim is to examine what level of signalling, signal-response and signal-dishonesty evolve at evolutionary equilibrium. To do this, we assume a monomorphic population, where all individuals have the same level of signalling as each other, given by *x*\*, as well as the same level of signal-response as each other, given by *y*\*, and the same level of signal-dishonesty as each other, given by *z\** (the lack of subscripts imply that the values are the same across the population).

We can then take a random (*focal*) individual and mutate its (and its identical-by-descent relatives’) level of signalling to a deviant value *x*, and ask how the fitness of the focal individual changes in response, in effect calculating 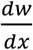. Analogously, we can mutate a random individual’s signal-response to *y* and calculate 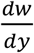. We can mutate a random individual’s signal-dishonesty to *z* and calculate 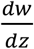. Evaluating these derivatives at equilibrium (*x*_*focal*_, *x*_*others*_, *x*_*pop*_ = *x*^*^, *y*_*focal*_, *y*_*others*_, *y*_*pop*_ = *y*^*^, *z*_*focal*_, *z*_*others*_, *z*_*pop*_ = *z*^*^), the equilibrium (ESS) levels of signalling (*x*\*), signal-response (*y*\*) and signal-dishonesty (*z*\*) can then be obtained, under the assumption of no genetic association (linkage disequilibrium) between traits, as the values for which 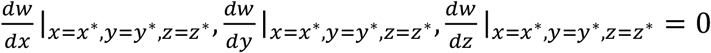 is satisfied (32).

We start by calculating 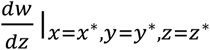 (mutating signal-dishonesty). We do so by expanding it using the chain rule

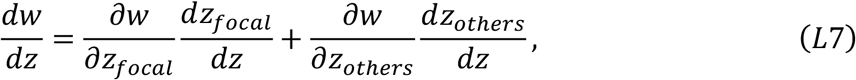

and then evaluating the derivatives on the right-hand side (3, 4, 32). Because the population is monomorphic, and because we are interested in the long-term evolutionary state of the population, we evaluate these derivatives at the point where *x*_*focal*_, *x*_*others*_, *x*_*pop*_ = *x*^*^, *y*_*focal*_, *y*_*others*_, *y*_*pop*_ = *y*^*^, *z*_*focal*_, *z*_*others*_, *z*_*pop*_ = *z*^*^. We trivially obtain

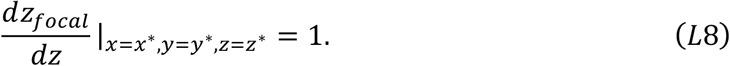

We obtain the equilibrium-evaluated partial derivatives 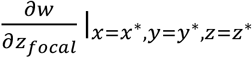 and 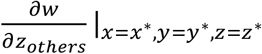 respectively as

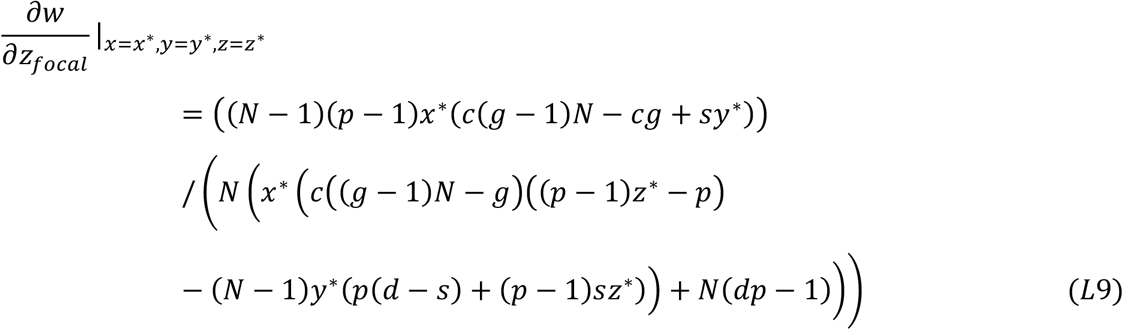

and

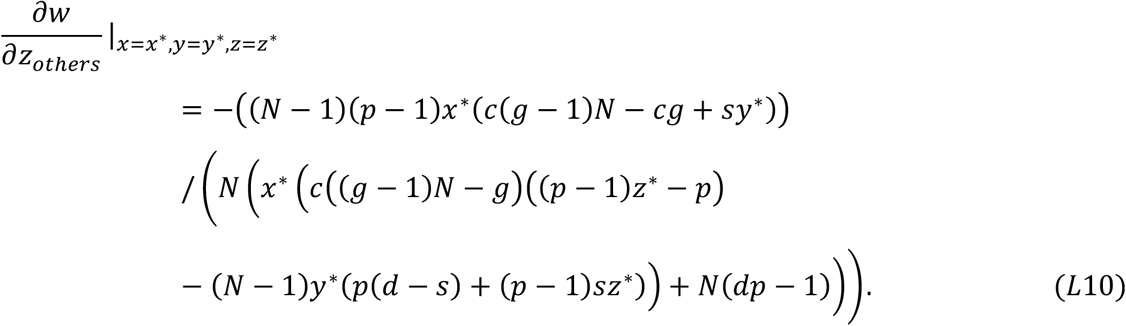

We obtain 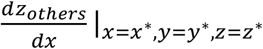 by noting that it is interpretable as a coefficient of relatedness, since it denotes the marginal (correlated) change in signal production by a social partner as a consequence of a change in signal production by an actor (3, 4). We can therefore write

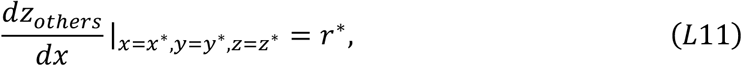

where *r*\* is the long-term (equilibrium) value of relatedness. We have already calculated *r*\* for our demographic assumptions (Appendix A; Equation A16), and we can substitute this into Equation L11 to obtain

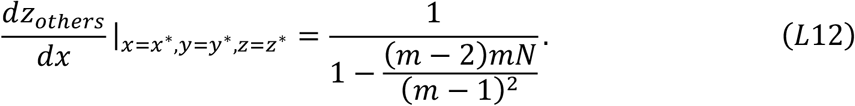

We can now substitute Equations L8, L9, L10 & L12 into Equation L7 to obtain an explicit expression for 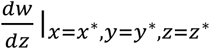. Setting 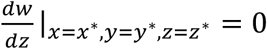 and simplifying, we obtain

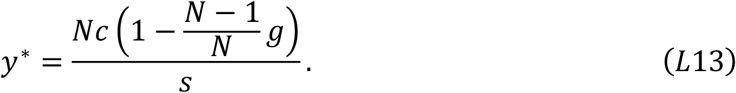

The equilibrium level of signal-dishonesty (*z*\*) corresponds to the value of *z*\* for which Equation L13 is satisfied (note that we haven’t obtained a specific value for *z*\* yet). To interpret Equation L13, note that 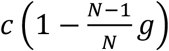 gives the cost of (dishonest) signalling to the actor. 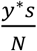 gives the cost of responding to a dishonest signal, borne by a competitor. Signal dishonesty is favoured to increase when it results in a greater cost to competitors than to self, meaning that, at an internal equilibrium, these costs to competitors and self are equalised, leading to 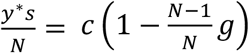, which with some slight rearrangement gives Equation L13.

Next, we calculate 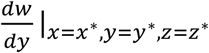 (mutating signal-response). We do so by expanding it using the chain rule

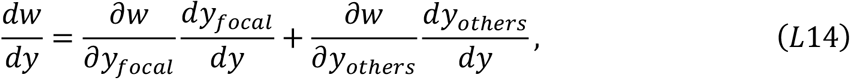

and then evaluating the derivatives on the right-hand side at equilibrium (3, 4, 32). We trivially obtain

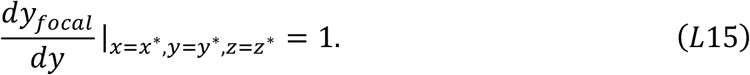

We obtain the equilibrium-evaluated partial derivatives 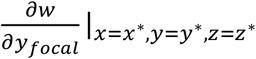 and 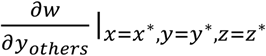 respectively as

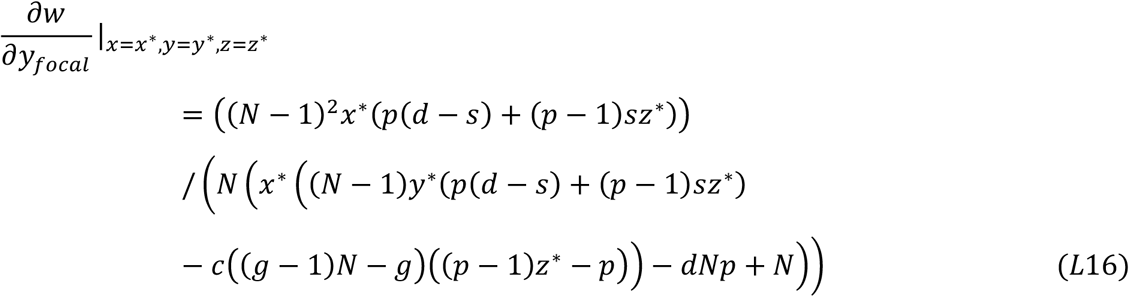

and

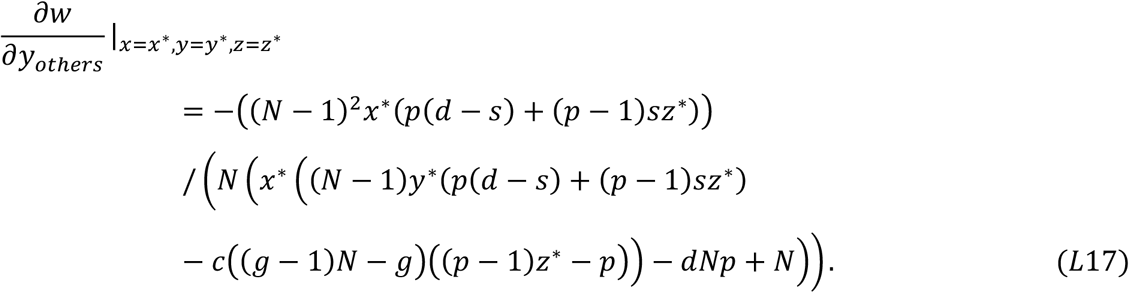

We obtain 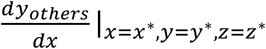 by noting that it is interpretable as a coefficient of relatedness, meaning we can write

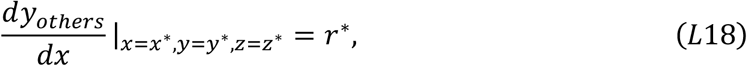

where *r*\* is the long-term (equilibrium) value of relatedness. Substituting in our explicit expression for *r*\* (Equation A16), we obtain

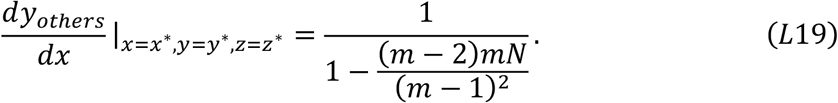

We can now substitute Equations L15, L16, L17 & L19 into Equation L14 to obtain an explicit expression for 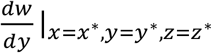. Setting 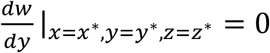 and simplifying, we obtain

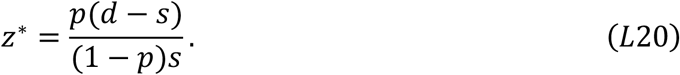

The equilibrium level of signal-response (*y*\*) corresponds to the value of *y*\* for which Equation L20 is satisfied (note that the specific value for *y*\* is provided in Equation L13). To interpret Equation L20, note that 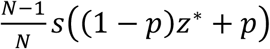 gives the cost of responding to a signal – specifically, it is the defence cost (*s*) weighted by the proportion of generations where this defence cost is paid 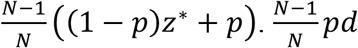 gives the benefit of responding to a signal – specifically, it is the benefit of a reduced attack cost (*d*), weighted by the proportion of generations where benefit can be accrued 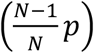. Signal response is favoured to increase when it results in a greater benefit than cost to the responder, meaning that, at an internal equilibrium, this benefit and cost are equalised, leading to *pd* = *s*((1 − *p*)*z* + *p*), which with some rearrangement gives Equation L20.

Finally, we calculate 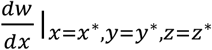 (mutating signal investment). We do so by expanding it using the chain rule 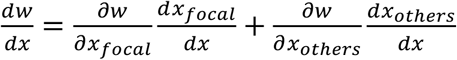 (Equation A7), and evaluating the derivatives on the right-hand side at equilibrium (3, 4, 32). We trivially obtain

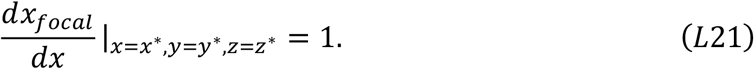

We obtain the equilibrium-evaluated partial derivatives 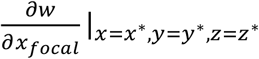 and 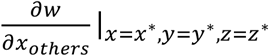 respectively as

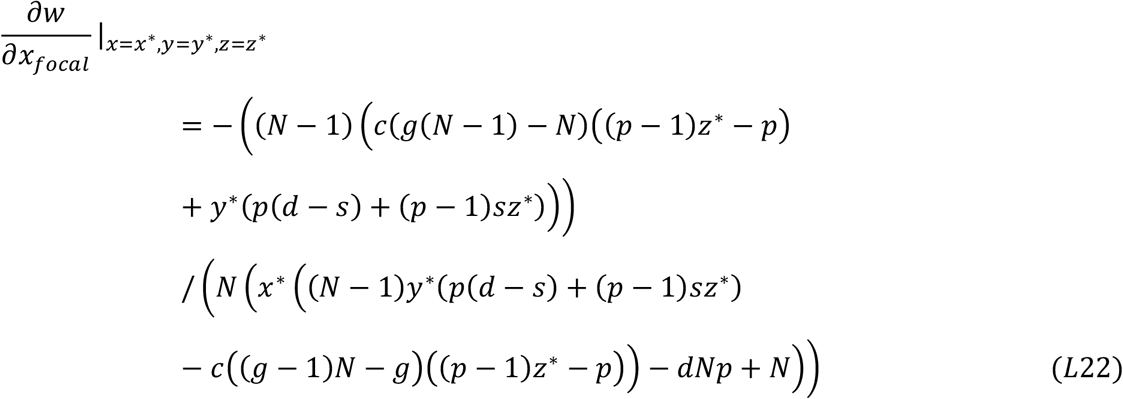

and

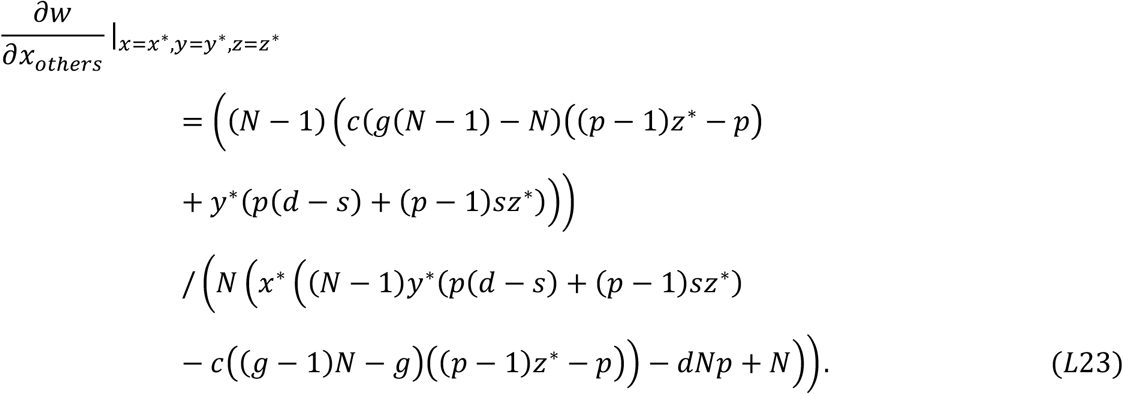

We obtain 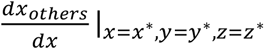 by noting that it is interpretable as a coefficient of relatedness, meaning we can write

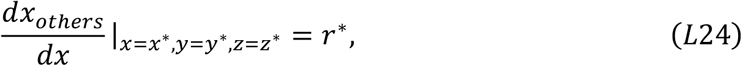

where *r*\* is the long-term (equilibrium) value of relatedness. Substituting in our explicit expression for *r*\* (Equation A16), we obtain

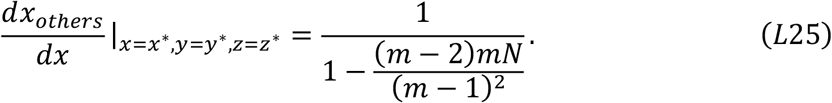

We can now substitute Equations L21, L22, L23 & L25 into Equation A7 to obtain an explicit expression for 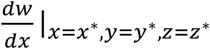. Instead of setting 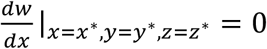 like we did for 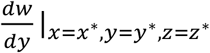 and 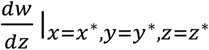, we set 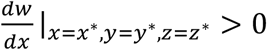 to see when signalling is favoured by natural selection (rather than when it is at an evolutionary equilibrium). We obtain

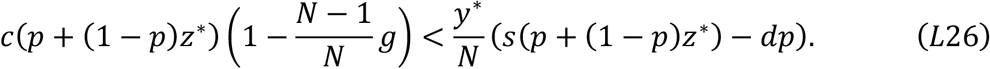

To interpret Equation L26, note that the left-hand side gives the overall cost of signalling to the actor, and the right-hand side gives the overall cost of signalling incurred by receivers. Signalling is then favoured in this lifecycle if hurts competitors more than oneself. Specifically, the cost of signalling to the actor (left-hand side) is obtained as the cost of signal production 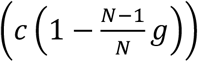 weighted by the proportion of generations where a signal is produced ((1 − *p*)*z*^*^ + *p*). The cost of signalling to receivers (right-hand side) is obtained as the cost of defence (*s/N*), where this is weighted by the proportion of generations where a signal is produced ((1 − *p*)*z*^*^ + *p*) as well as by the response to the signal (*y*\*), minus the saved cost of attack (d/N), where this is weighted by the proportion of generations where a herbivore is present (p) as well as by the response to the signal (*y*\*).

We can obtain explicit values for *y*\* and *z*\*. These explicit values will often be the values of *y*\* and *z*\* specified by Equations L13 & L20. However, *y*\* and *z*\* must lie in the range 0 to 1, but the value of *y*\* or *z*\* specified by Equations L13 & L20 may sometimes go above 1. In such cases, the true (obtained) value for *y*\* or *z*\* will be 1. Similarly, the value of *y*\* or *z*\* specified by Equations L13 & L20 may sometimes fall below zero. In such cases, the true (obtained) value for *y*\* or *z*\* will be 0. Substituting these explicit values for *y*\* and *z*\* into Equation L26, we find that Equation L26 is never satisfied, meaning signalling is never favoured (*x*\*=0) (see supplementary *Matlab* file for a numerical demonstration of this).

This recovers the result of Model A. Therefore, the result of Model A, that signalling does not evolve, still holds, after generalising the model to allow dishonest signalling and submaximal signal-response. The reason why this previous result holds is that, in this generalised model, signals either evolve to be relatively more dishonest or honest. Dishonest signals harm competitors by inducing them to defend themselves against herbivores that aren’t even there, giving the signaller a transient advantage, but inducing competitors to ignore the signal. Ultimately, the signal is ignored, meaning costly signal production is disfavoured. Conversely, honest signals help competitors by warning them about herbivores, giving the signaller a competitive disadvantage, disfavouring signal production. Therefore, irrespective of how honest or dishonest the signal evolves to be, signal production is ultimately disfavoured in this lifecycle.

We note that ‘opening’ this model, detaching relatedness from demographic parameters (*m* and *N*) so that it is a parameter in its own right, does not affect model outcomes. To appreciate this, note that calculating 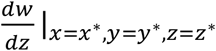 with Equation L11 rather than Equation L12 (i.e., treating r* as a parameter rather than a function of demographic parameters) does not lead to a different expression for 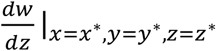. Similarly, calculating 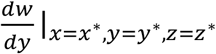 with Equation L18 rather than Equation L19 does not lead to a different expression for 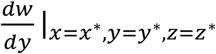. Calculating 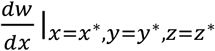 with Equation L24 rather than J25 does not lead to a different expression 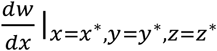. This means that the results of the model (Equations L13, L20 & L26) are unchanged – relatedness does not affect model outcomes in this lifecycle.

## Appendix M

### Model M: signalling; open & closed; population regulation after migration; individuals may signal dishonestly and not respond to signals

In this appendix, we alter Model L so that population regulation occurs after migration rather than before. We examine both ‘closed’ (emergent relatedness) and ‘open’ (parameterised relatedness) versions of the model, starting with the open version. We found in Appendix D that signalling can evolve in the open version of this lifecycle (with population regulation occurring after migration). However, this result was obtained on the assumptions that signals are honest and individuals respond maximally to signals. In this appendix, we see if this result (that signalling can evolve) still holds when signal-dishonesty and signal-response can evolve.

Our assumption that population regulation occurs after migration means that the fitness of the focal individual is given by Equation C1 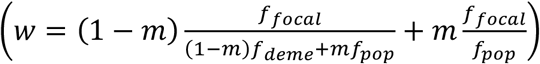, where explicit expressions for the fecundity (*f*_*pop*_, *f*_*deme*_, *f*_*pop*_) functions are provided in Equations L3, L5 & L6 (7, 12, 13).

Our aim is to examine what level of signalling, signal-response and signal-dishonesty evolve at evolutionary equilibrium, as we did in Appendix L. We do so by first calculating 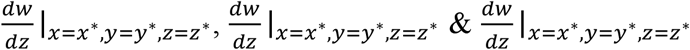 by expanding them using the chain rule (Equations L7, L14 & A7), and evaluating the associated partial derivatives (3, 4, 32). We obtain the following partial derivatives:

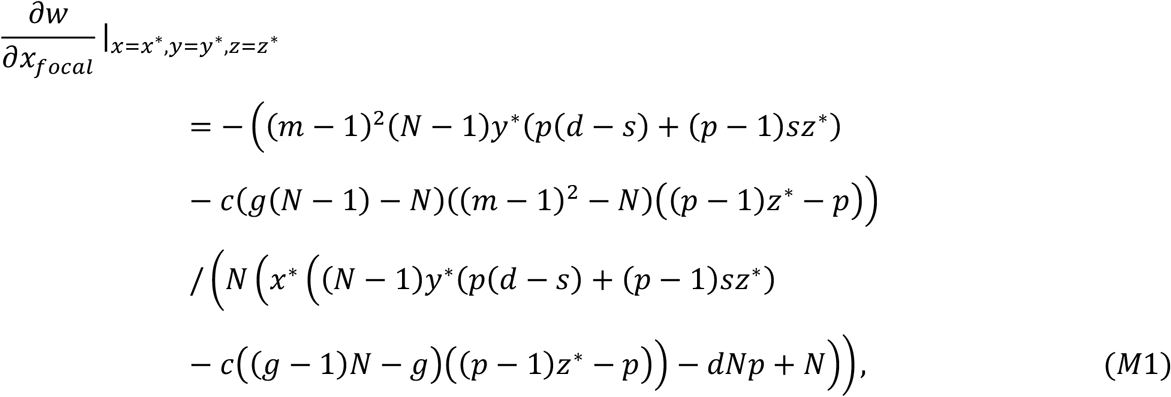

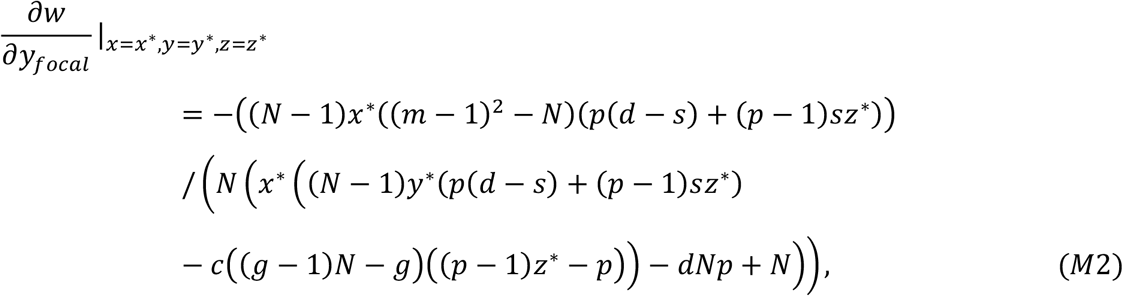

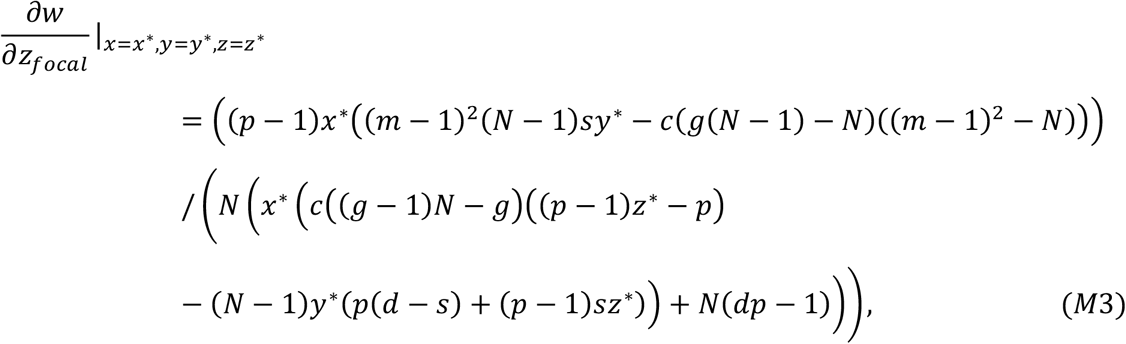

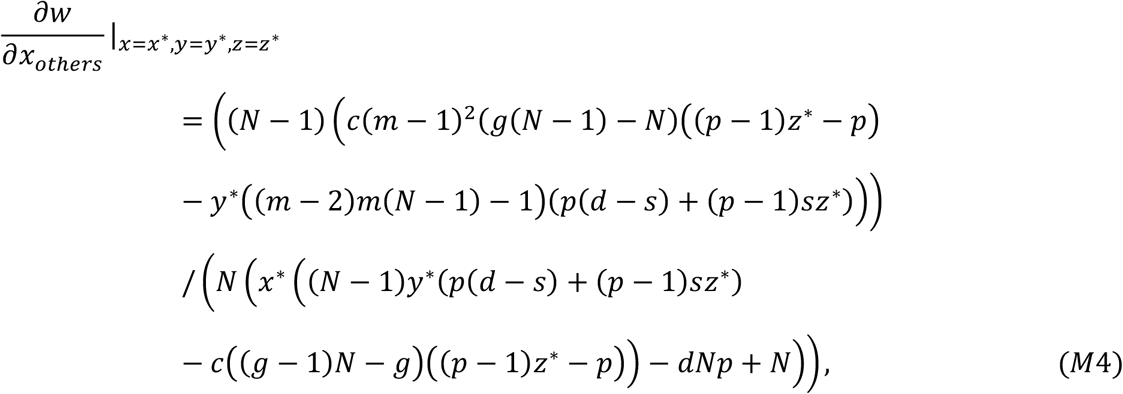

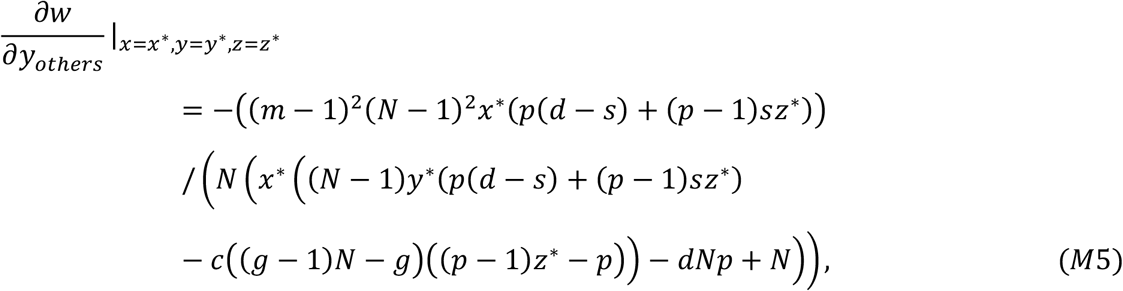

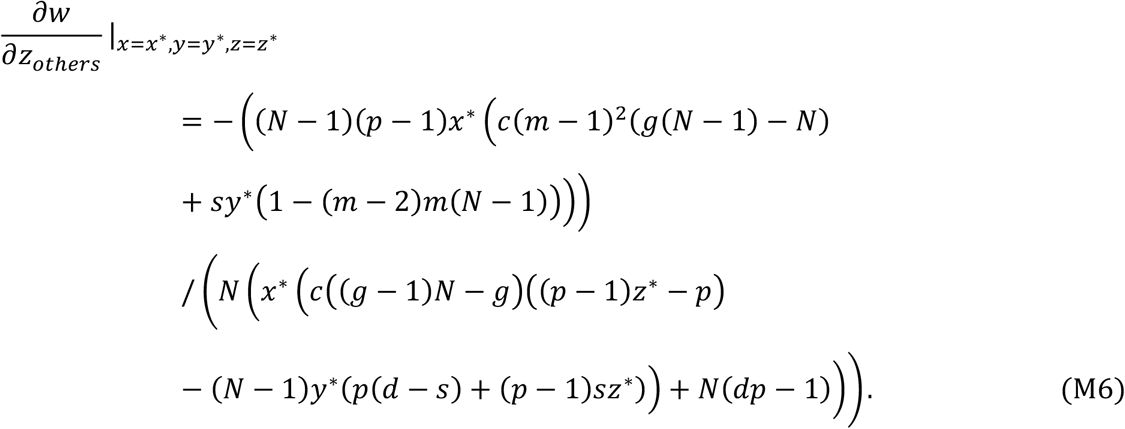

We can now substitute Equations L21, L24, M1 & M4 into Equation A7 to obtain an explicit expression for 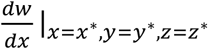. Analogously, we can substitute Equations L15, L18, M2 & M5 into Equation L14 to obtain an explicit expression for 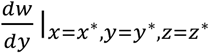. We can substitute Equations L8, L11, M3 & M6 into Equation L7 to obtain an explicit expression for 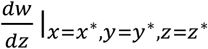.

We find that the explicit value for 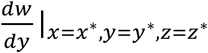 is the same in this model as it was for Model L (Equation L20). Setting 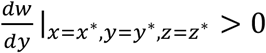 and simplifying, we obtain

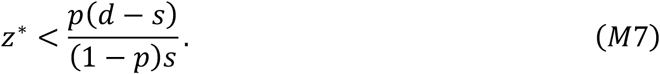

Setting 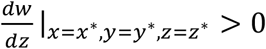 and simplifying, we obtain

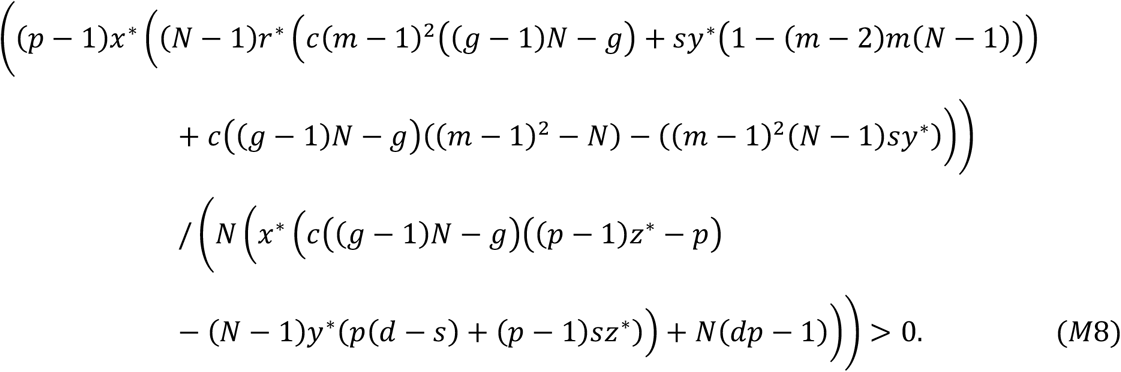

Setting 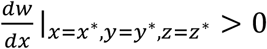 and simplifying, we obtain

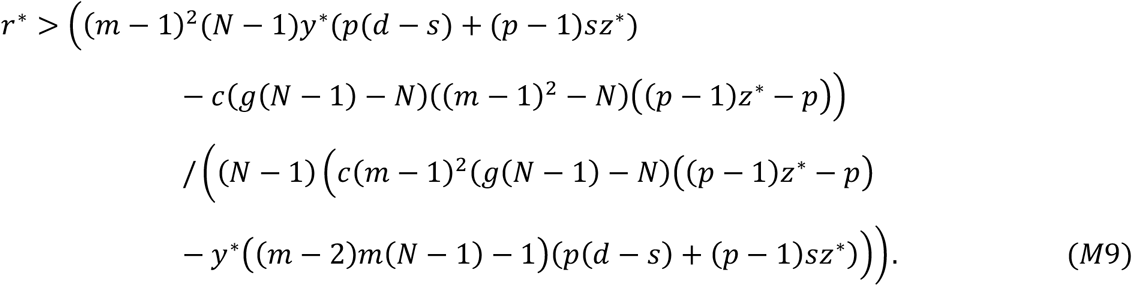

Signal-response is favoured if Equation M7 is satisfied. Conversely, signal-response is disfavoured if 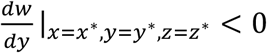, and at an internal ESS if 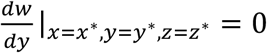. Signal-dishonesty is favoured if Equation M8 is satisfied. Conversely, signal-dishonesty is disfavoured if 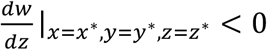, and at an internal ESS if 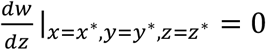. Signalling is favoured Equation M9 is satisfied. Conversely, signalling is disfavoured if 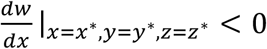, and at an internal ESS if 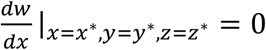.

Equations M7, M8 & M9 describe how selection acts on each of the three traits (response, dishonesty & signalling). We cannot solve this model analytically, so we use numerical methods to elucidate what trait values obtain in the evolutionary long term (see supplementary *Matlab* file). Our approach assumes that there is no genetic correlation (linkage disequilibrium) between the traits (32). Specifically, we assume some initial values for *x, y & z* and use them to calculate the selection differential for each of the three traits 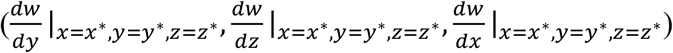. If the selection differential is positive, we add on a very small increment to the previous trait value; if the selection differential is negative, we subtract a very small increment; if the selection differential is zero, we do not add or subtract an increment. We then use the new trait values to calculate the selection differentials, and repeat the process until trait values have stopped changing. We find that the resulting trait values are independent of initial trait values and of the size of the increment, as long as the increment is sufficiently small. The resulting trait values correspond to the long-term trait values: *x*, y* & z\**.

We find that, in this lifecycle, signals evolve to be honest and responded-to at equilibrium, and that signalling evolves in the same region of parameter space that it did in Model D, which assumed the same lifecycle but forced signals to be honest and responded-to.

Specifically, we find that, when the right-hand side of Equation D1 (which gives an explicit value for 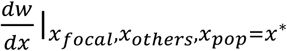 for Model D) is greater than zero, signalling is favoured in the present model (*x*\*=1), and the signals are favoured to be honest (*z*\*=0) and responded-to (*y*\*=1). Conversely, when the right-hand side of Equation D1 is less than zero, signalling is not favoured in the present model (*x*\*=0), and any signals that are (transiently) present in the population are favoured to be dishonest (*z*\*=1) and ignored (*y*\*=0). The reader may wish to consult Fig. S1, which can be found in Appendix D – signalling, signal honesty and signal response are favoured above the lines.

As can be seen by inspecting Equation D1, a necessary requirement for the evolution of honest signalling is this is that there is sufficient relatedness and migration. The reason for this is that high relatedness leads to social interactions with kin, and high migration leads to competition with non-kin, and these combined conditions are necessary for the evolution of helping behaviours, such as honestly signalling the presence of a herbivore.

We can ‘close’ this model by substituting an explicit expression for relatedness (*r*\*) into our results conditions. Specifically, we substitute Equation A16 into Equations M8 and M9 and use the same numerical method to find long-term trait values. Doing so, we find that signalling can no longer evolve. This recovers the result of Model C, which made the same demographic assumptions (same lifecycle) but didn’t allow signal-dishonesty or signal-response to evolve.

## Appendix N

### Model N: monitoring; open & closed; population regulation before migration; individuals may not respond to signals

In plants, information about the presence of herbivores is often transmitted between individuals through common (shared) mycorrhizal networks. The mycorrhizal network is an evolutionary agent (individual) in itself, which raises the possibility that the mycorrhizal network might ‘monitor’ a plant, to find out when it is being attacked by a herbivore, and then signal this information to the other plants on the network. We examine this possibility in this Appendix.

This model differs from previous models in that it is a coevolutionary model (*i*.*e*., it tracks evolution in two rather than one species), and that the signaller is the fungus rather than the plant. We examine both ‘closed’ (emergent relatedness) and ‘open’ (parameterised relatedness) versions of the model, starting with the closed version.

We assume an infinite populations of plants, as well as an infinite population of fungi. We assume for simplicity that all individuals (plants and fungi) are haploid and asexual. The plant and fungi populations are split into demes, where each deme has *N* plants and 1 fungus, where the *N* plants are connected to the fungus (common mycorrhizal network) and therefore to each other. We focus our analysis on a ‘focal plant’ and a ‘focal fungus’, drawn at random from the population.

Each generation, with probability *p*, the population is attacked by herbivores (with probability 1-*p*, the population is not attacked). In generations where the population is attacked, a random plant on each deme is initially attacked by a herbivore, and suffers a fecundity cost of *d*. Given that there are *N* plants on each deme, each plant has a 1/*N* chance of being initially attacked by the herbivore. The fungus may then invest into ‘monitoring & signalling’ to learn that the initially-attacked individual has been attacked, and warn the other *N*-1 plants on the deme of this. Specifically, the ‘focal fungus’ invests an amount 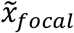, into monitoring & signalling (the tilde over the *x* denotes fungal rather than plant behaviour). Monitoring & signalling investment incurs a fecundity cost that scales linearly with *c*. Therefore, the focal fungus pays a monitoring & signalling cost of 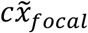.

The signal produced by the fungus is transferred to (and recognised by) the other *N*-1 individuals on the deme. The signal warns the other plants that a herbivore is in the vicinity. Consequently, the other plants can prepare for being attacked, and defend themselves, meaning they can reduce their fecundity cost of attack. Plants may choose the extent to which they actually respond to the signal, preparing for herbivore-attack, rather than ignoring the signal. Preparation for herbivore attack (defence) incurs a fecundity cost that scales linearly with *s*. We assume that there is no genetic correlation between fungal and plant genotypes (i.e., no partner choice), which means that the focal plant’s mycorrhizal network (fungus) invests 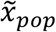 into monitoring & signalling, where the *pop* subscript denotes the (fungal) population average investment. We denote the extent of signal-response by the focal individual by *y*_*focal*_. This means that, when the focal individual is not the initially-attacked individual, it will ultimately suffer a cost of attack that is given by 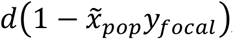, and a defence cost that is given by 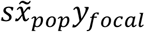. We assume that preparing for herbivore attack is less costly than the fecundity cost of being attacked (*d*>*s*), which ensures that individuals are favoured to defend themselves against herbivores.

In generations where the population is not attacked by herbivores (such generations occur with probability 1-*p*), fungal individuals are given the opportunity to produce the signal of herbivore-presence even though no herbivore is present. In such cases, any signal produced is ‘dishonest’. The focal fungus invests 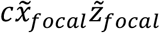, into signal production, where 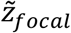, denotes dishonesty, with 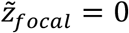 corresponding to no dishonest signal production, and 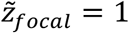 corresponding to the same investment in signal production irrespective of whether the herbivore is present or not. Dishonest signalling means that the focal plant suffers a cost of preparation for herbivore attack that is given by 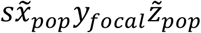.

We assume that fungi gain a benefit by being connected to ‘fitter’ (more fecund) plant partners, because such partners will be better trade partners (*i*.*e*., they will be in a position where they can acquire and transfer carbon to their fungal mutualist). Specifically, we assume that the focal fungus gains a benefit of *bf*_*partner*_, where *f*_*partner*_ gives the average fecundity of the plants connected to the focal fungus.

After fecundity effects have been incurred by all individuals, each individual produces a large number of offspring (juvenile haploid clones) in proportion to their fecundity. Population regulation and migration for the plant population then occurs as follows. A random sample of *N* juvenile plants are chosen, for each deme, to survive and form the next adult population (local population regulation). After population regulation, a proportion of the (new) adult plant population, *m*, migrate to a different deme, randomly chosen for each individual. The remaining proportion, *1-m*, remain on their local deme.

Population regulation and migration for the fungal population occurs as follows. Each member of the juvenile population migrates (with certainty) to a different deme, randomly chosen for each individual. A random juvenile individual is then chosen, for each deme, to survive and be the ‘adult’ fungus (mycorrhizal network) for that deme in the next generation. Note that the assumption of certain migration followed by population regulation for the fungal population is taken to ensure that population regulation occurs effectively at the global rather than local level. Local population regulation would prevent evolution from occurring in this case, since there is just one genotype per deme, meaning natural selection would be ‘choosing’ between individuals with the same genotype. This lifecycle then iterates over many generations until an evolutionary end point is reached.

We denote the population average plant investment into signal response by *y*_*pop*_. We denote the average investment into signal response on the focal plant’s deme (incorporating its own signal response) by *y*_*deme*_; this can be written in terms of *y*_*focal*_ and *y*_*others*_ as 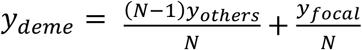 (Equation L1). We assume that the baseline fecundity is 1, that 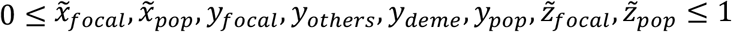, that *c* ≤ 1 and that *d* + *s* ≤ 1; these assumptions ensure that fecundity never falls below zero. The number of juvenile offspring produced by the focal plant (*f*_*focal*_) and focal fungus 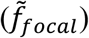 are then, respectively, proportional to

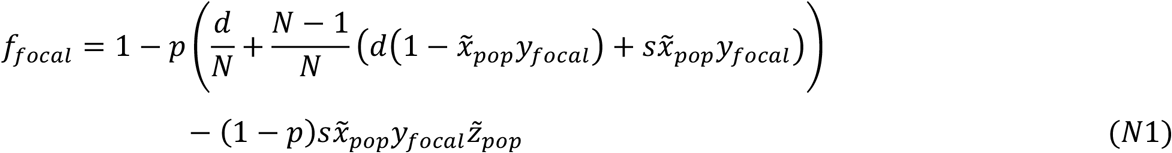

and

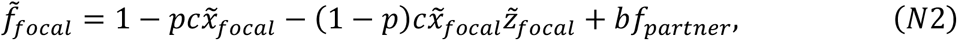

where *f*_*partner*_ denotes the average fecundity of plants connected to the focal fungus, and is proportional to

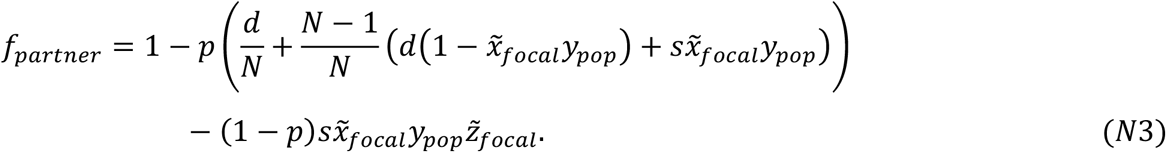

We can make sense of the right-hand side of Equation N1 as follows. Baseline plant fecundity is one (left-hand term). With probability *p*, a herbivore attacks. Given herbivore-attack, with probability 1/*N*, the focal plant is attacked first, leading to an additively applied cost of attack (*d*). Given herbivore-attack, with probability 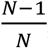, the focal plant is not attacked first, and can receive a signal of herbivore presence from the fungus it is connected to, leading to additively applied costs of attack 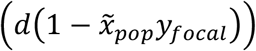 and defence 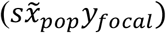. With probability 1-*p*, a herbivore does not attack. Given herbivore-absence, the fungus connected to the focal plant is given the opportunity to dishonestly signal herbivore-presence, meaning the focal plant suffers an additively applied cost of defence 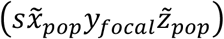. Equation N3 is obtained *mutatis mutandis*.

We can make sense of the right-hand side of Equation N2 as follows. Baseline fungus fecundity is one (left-hand term). With probability *p*, a herbivore attacks, meaning the focal fungus incurs an additively applied cost of honest signal production 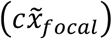. With probability 1-*p*, a herbivore does not attack, meaning the focal fungus incurs an additively applied cost of dishonest signal production 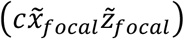. The focal fungus also gains an additively applied benefit of being connected to high-quality (fecund) trade partners (*bf*_*partner*_).

The number of juvenile offspring produced by a plant drawn randomly from the focal plant’s deme (including the focal individual) is proportional to

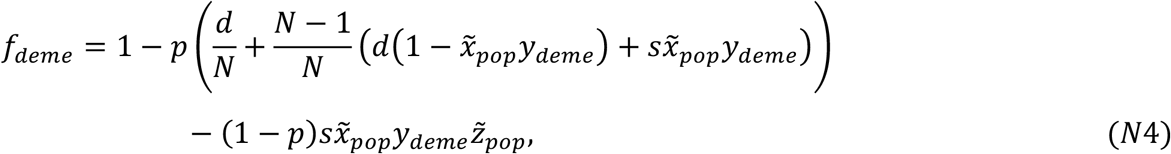

which we can rewrite using Equation L1 (getting rid of the superfluous *y*_*deme*_ parameter) as

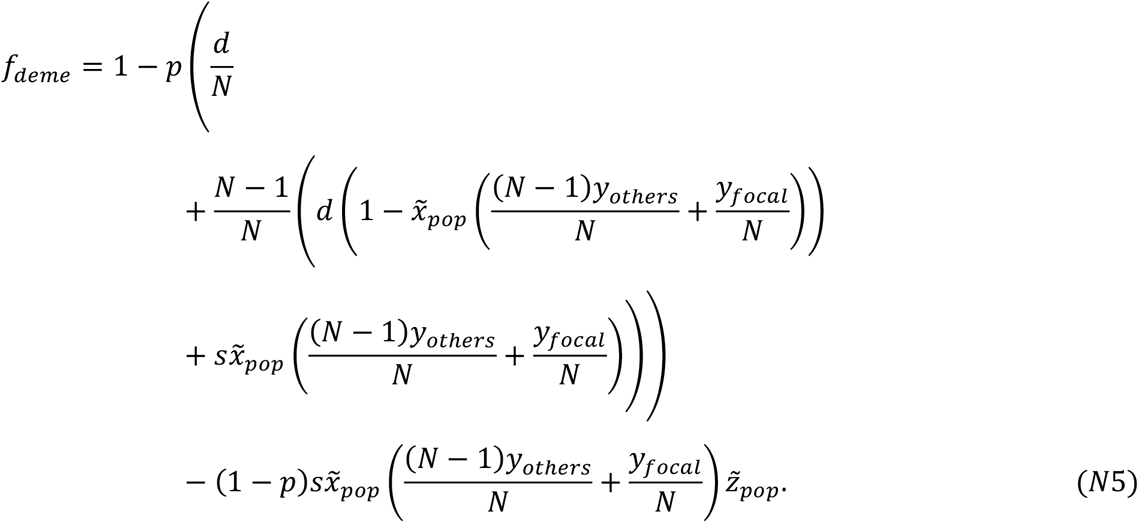

The number of juvenile offspring produced by a plant drawn randomly from the population is proportional to

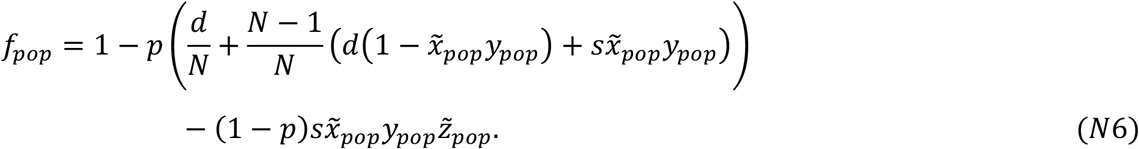

The number of juvenile offspring produced by a fungus drawn randomly from the population is proportional to

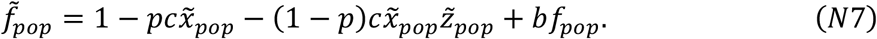

The fitness of the focal plant is then given by Equation A6 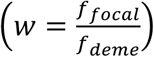, where explicit expressions for the fecundity (*f*_*focal*_, *f*_*deme*_) functions are provided in Equations N1 & N5 (7, 12, 13). The fitness of the focal fungus is given by

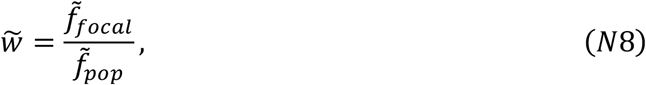

where explicit expressions for the fecundity 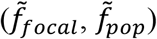 functions are provided in Equations N2 & N7 (we also need Equation N6 to evaluate the *f*_*pop*_ term in Equation N7).

Our aim is to examine what level of fungal monitoring & signalling, plant signal-response, and fungal signal-dishonesty evolve at evolutionary equilibrium. To do this, we assume a monomorphic fungus population, where all fungi have the same level of monitoring & signalling as each other, given by 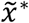, as well as the same level of signal-dishonesty as each other, given by 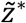. We additionally assume a monomorphic plant population, where all plants have the same level of signal-response as each other, given by *y*\*.

We can then take a random (focal) fungus and mutate its (and its identical-by-descent relatives’) level of signalling to a deviant value *x*, and ask how the fitness of the focal fungus changes in response, in effect calculating 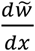 (to reiterate, the tilde over the *w* is used to refer to fungal rather than plant fitness). Analogously, we can mutate a random fungus’ signal-dishonesty to *z* and calculate 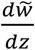. Additionally, we can mutate a random plant’s signal-response to *y* and calculate 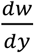. Evaluating these derivatives at equilibrium 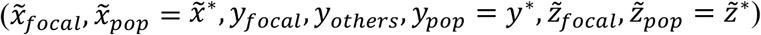, the equilibrium (ESS) levels of signalling 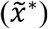, signal-response (*y*\*) and signal-dishonesty 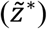 can then be obtained, under the assumption of no genetic association (linkage disequilibrium) between traits, as the values for which 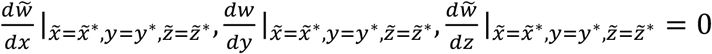 is satisfied (32–34).

We start by calculating 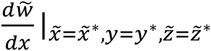 (mutating signal-dishonesty). We do so by expanding it using the chain rule

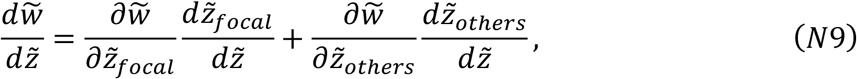

and then evaluating the derivatives on the right-hand side at equilibrium (3, 4, 32). Because the plant and fungus populations are monomorphic, and because we are interested in the long-term evolutionary state of the population, we evaluate these derivatives at the point where 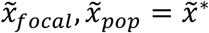, *y*_*focal*_, *y*_*others*_, 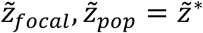. We trivially obtain

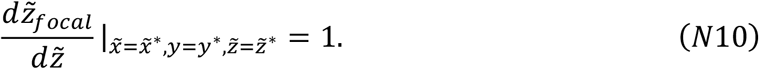

We obtain the equilibrium-evaluated partial derivatives 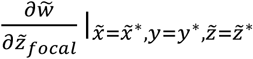 and 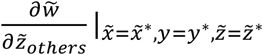 respectively as

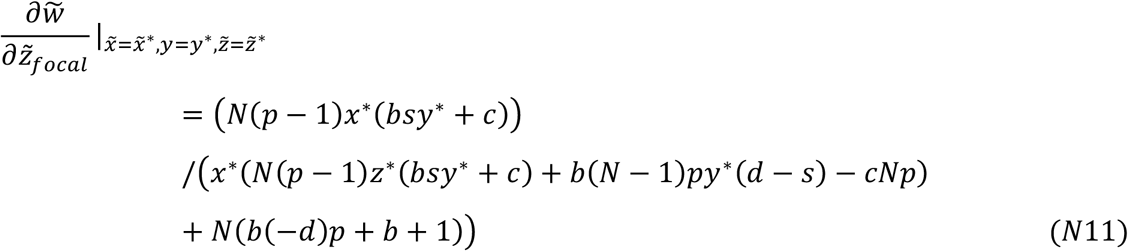

and

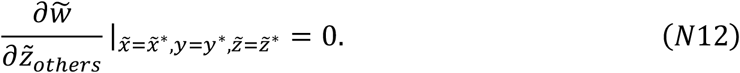

Note that, because 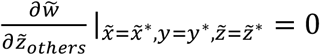 (Equation N12), this gets rid of the 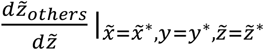 partial derivate in Equation N9, which means we don’t need to evaluate 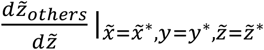 in order to evaluate the full derivative 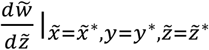.

We can now substitute Equations N10, N11 & N12 into Equation N9 to obtain an explicit expression for 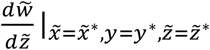. Setting 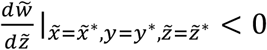 and simplifying, we obtain

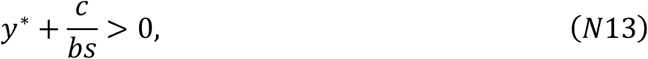

which we can see is satisfied (as long as *y*\*>0 or *c*>0), meaning signal dishonesty is disfavoured, resulting in an equilibrium signal dishonesty of

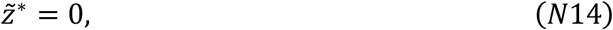

corresponding to completely honest signals. The reason for this result is that, in this model, the fungi are the signallers, and they can never gain a fitness advantage from signalling dishonestly. Dishonest signalling incurs an energetic cost to the fungus (proportional to *c*). Furthermore, it reduces the fecundity of the fungus’ plant partners, since they invest in herbivore defence when no herbivore is present (proportional to *s*), which in turn reduces the fitness of the fungus as a consequence of having lower quality trade partners (proportional to *b*). Signalling dishonestly is therefore always bad for the signaller (fungus), meaning fungi are favoured to increase the honesty of their signals.

Next, we calculate 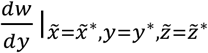 (mutating signal-response). We do so by expanding it using the chain rule 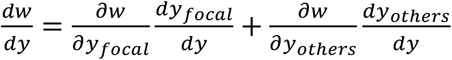 (Equation L14), and then evaluating the derivatives on the right-hand side at equilibrium (3, 4, 32). We trivially obtain

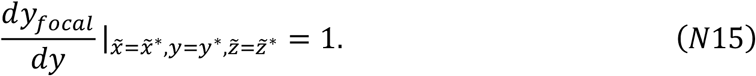

We obtain the equilibrium-evaluated partial derivatives 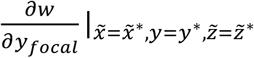 and 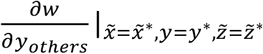 respectively as

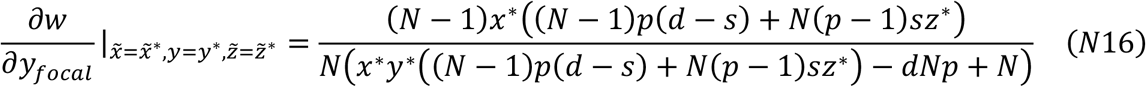

and

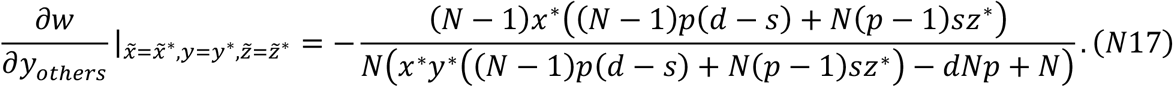

We obtain 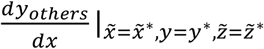 by noting that it is interpretable as a coefficient of relatedness (between plant individuals), meaning we can write

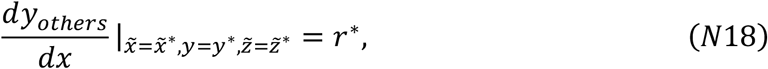

where *r*\* is the long-term (equilibrium) value of plant relatedness. Substituting in our explicit expression for *r*\* (Equation A16), we obtain

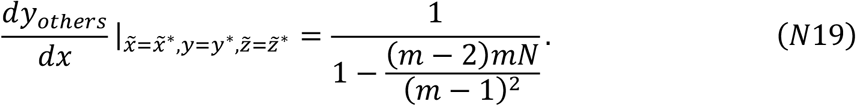

We can now substitute Equations N15, N16, N17 & N19 into Equation L14 to obtain an explicit expression for 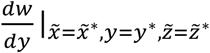. Setting 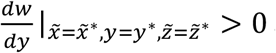 and simplifying, we obtain

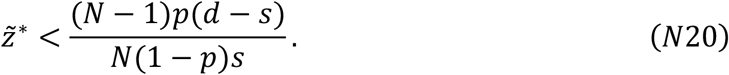

Signal-response (*y*\*) is favoured to increase if Equation N20 is satisfied. To interpret Equation N20, note that 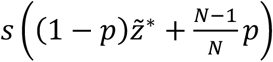 gives the cost to a plant of responding to a signal – specifically, it is the defence cost (*s*) weighted by the proportion of generations where this defence cost is paid 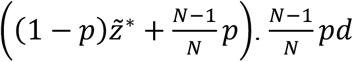 gives the benefit to a plant of responding to a signal – specifically, it is the benefit of a reduced attack cost (*d*), weighted by the proportion of generations where benefit can be accrued 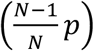. Signal response is favoured to increase when it results in a greater benefit than cost to the responder (plant), leading to 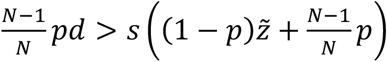, which with some rearrangement gives Equation N20.

Note that the right-hand side of Equation N20, which gives the threshold signal dishonesty under which signal-response is favoured, is almost the same as the right-hand side of Equation L20. The difference is that the right-hand side of Equation N20 is a factor 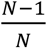 greater than the right-hand side of Equation L20. The reason for this difference is that, in the previous model where plants are signallers (Model L), each plant has a generational probability 1/*N* of being ‘chosen’ to be a potential signaller rather than a potential signal receiver. In the present model where fungi are signallers, in generations where a herbivore is absent and the fungus signals dishonestly, all plants on the deme receive this signal (*i*.*e*., there is no ‘chosen’ plant that does not receive the signal). This slightly increased probability of receiving a signal in the present model, relative to Model L, is the reason why the 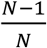 factor features in Equation N20.

By substituting our explicit value for signal dishonesty (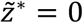; Equation N14) into our condition for signal-response being favoured (Equation N20), we find that Equation N20 is satisfied (as long as *p*>0 & *d*>0). This means that signal-response is favoured, leading to

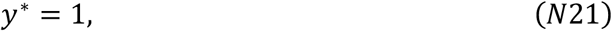

corresponding to maximal response of plants to fungal signals. The reason why plants are favoured to respond maximally to fungal signals in this model is that, at equilibrium, fungal signals are honest, meaning they reliably indicate that a herbivore is present (Equation N14). It is always in the plants’ interests to respond to such signals and defend themselves against the impending herbivore attack.

Finally, we calculate 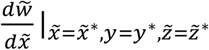 (mutating signal investment). We do so by expanding it using the chain rule

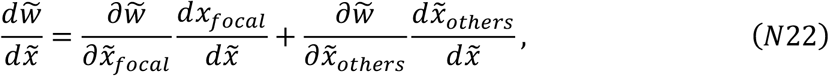

and evaluating the derivatives on the right-hand side at equilibrium (3, 4, 32). We trivially obtain

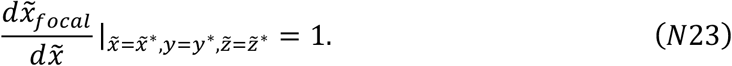

We obtain the equilibrium-evaluated partial derivatives 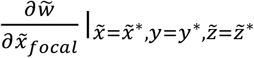 and 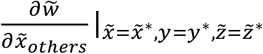 respectively as

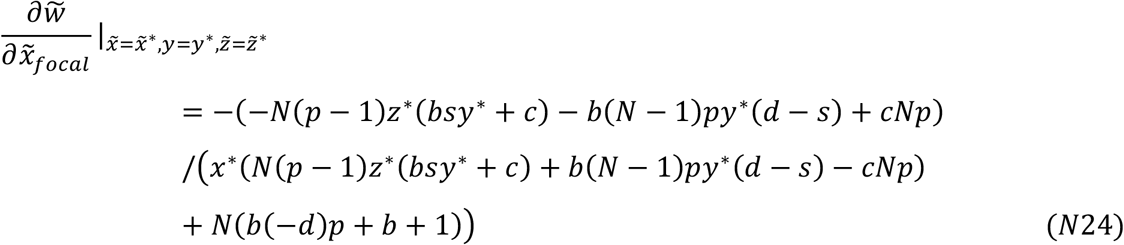

and

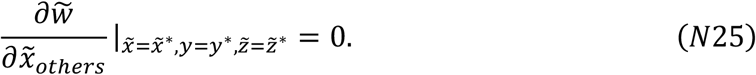

Note that, because 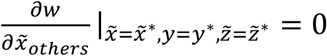 (Equation N24), this gets rid of the 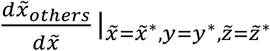 partial derivate in Equation N21, which means we don’t need to evaluate 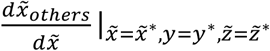 in order to evaluate the full derivative 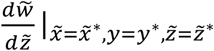.

We can now substitute Equations N23, N24 & N25 into Equation N22 to obtain an explicit expression for 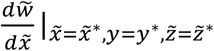. Setting 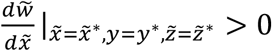, substituting in our explicit values for *y\** (Equation N21) and 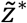 (Equation N14), and simplifying, we obtain

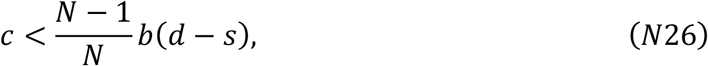

which is a condition for when fungi are favoured to signal the presence of a herbivore to the plants that they are connected to. Signalling is favoured when the benefit of signalling to the signaller (fungus), given by the right-hand side of Equation N26, is greater than the cost of signalling to the signaller (fungus), given by the left-hand side of Equation N26. The benefit of signalling to the signaller (fungus) is obtained as the increase in plant fecundity (fitness) as a consequence of responding to the fungal signal and upregulating herbivore defence (*d*–*s*), weighted by the fraction of the plant population who can respond to the fungal signal and therefore obtain this increase in fecundity 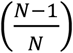, weighted by the benefit to the fungus of having higher-quality trade partners (*b*). The cost of signalling to the signaller (fungus) is simply given by the cost of signal production (*c*).

We note that ‘opening’ this model, detaching relatedness from demographic parameters (*m* and *N*) so that it is a parameter in its own right, does not affect model outcomes. To appreciate this, note that calculating 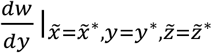 with Equation N18 rather than Equation N19 (i.e., treating *r*\* as a parameter rather than a function of demographic parameters) does not lead to a different expression for 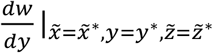. Relatedness does not feature in the expressions for 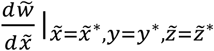 and 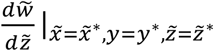. This means that the results of the model (Equations N14, N21 & N26) are unchanged – relatedness does not affect model outcomes in this lifecycle.

## Appendix O

### Supplementary discussion about ‘competition’ in our models

Our result that signalling between plants tends to be disfavoured hinges on ‘competition’. Signalling tends to be disfavoured because signalling provides a benefit to individuals who are in direct competition with the signaller.

Our models are island models, which means that competition arises because individuals have lots of offspring, but only a limited number of these offspring (*N*) are given spots on the patch to grow into adults. There is therefore local competition (=density dependence). This is the standard way of modelling competition in population genetics (*e*.*g*., 12) and social evolution (*e*.*g*., 4).

Mathematically, competitive effects are captured in our fitness functions, *e*.*g*. 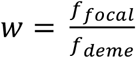 (Equation A6) and 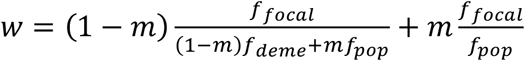 (Equation C1). These equations divide personal fecundity through by the average fecundity of competitors (for that lifecycle). The division guarantees that fitness is a ‘relative’ rather than ‘absolute’ measure, such that fitness is reduced if the fecundity of competitors is increased. We note that an alternative mathematical approach would have been to write fitness as a sequence of additive terms, with one of these terms accounting for competition (*e*.*g*., the mathematical approach taken by Scott et al. (31)). However, the two approaches are mathematically equivalent. In other words, dividing through by the average fecundity of competitors (as we do here) means that competition is accounted for in the maths, without explicitly having a term for competition.

An alternative way of conceptualising competition could have been to assume that it directly affects fecundity (rather than arising as a consequence of lots of juveniles competing for spots on demes to grow into adults). In this scenario, each individual would initially have their fecundity modulated by social interactions and herbivory as usual, resulting in a ‘primary’ fecundity. But then each individual would need to have their fecundity modulated again, to ensure that average fecundity in the population and average fecundity amongst competitors on a deme are equal to one (this guarantees that the population remains constant in size), to obtain a ‘secondary’ fecundity that is equivalent to ‘fitness’. Secondary fecundity (fitness) would be obtained by dividing primary fecundity through by the average primary fecundity of competitors. This is exactly what we did in our models. Consequently, modelling competition as directly impacting fecundity is mathematically equivalent to the way we have modelled competition in our models.

More generally, ‘competition’ can be envisaged as a displacement that occurs when a population has a total reproductive value (fitness) that exceeds resources. It doesn’t matter whether competition exerts its effect by modulating survival or fecundity; what matters is that it brings the average fitness within the population (and within population classes) down to be in line with the available resources (5). In populations of constant size such as ours, this means that average population fitness is brought to 1.

## Notes

### Competing Interest Statement

The authors have declared no competing interest.

### Summary of Updates

Final manuscript and supplementary information for "The evolution of signalling and monitoring in plant-fungal networks" (Scott, Kiers & West).

